# The adaptive transcriptional response of pathogenic *Leptospira* to peroxide reveals new defenses against infection-related oxidative stress

**DOI:** 10.1101/2020.04.03.015982

**Authors:** Crispin Zavala-Alvarado, Odile Sismeiro, Rachel Legendre, Hugo Varet, Giovanni Bussotti, Jan Bayram, Samuel Garcia Huete, Guillaume Rey, Jean-Yves Coppée, Mathieu Picardeau, Nadia Benaroudj

## Abstract

Pathogenic *Leptospira* spp. are the causative agents of the waterborne zoonotic disease leptospirosis. During infection, *Leptospira* are confronted with dramatic adverse environmental changes such as deadly reactive oxygen species (ROS). Withstanding ROS produced by the host innate immunity is an important strategy evolved by pathogenic *Leptospira* for persisting in and colonizing hosts. In *L. interrogans*, genes encoding defenses against ROS are repressed by the peroxide stress regulator, PerR. In this study, RNA sequencing was performed to characterize both the *L. interrogans* adaptive response to low and high concentrations of hydrogen peroxide and the PerR regulon. We showed that *Leptospira* solicit three main peroxidase machineries (catalase, cytochrome C peroxidase and peroxiredoxin) and heme to detoxify oxidants produced during a peroxide stress. In addition, canonical molecular chaperones of the heat shock response and DNA repair proteins from the SOS response were required for *Leptospira* recovering from oxidative damages. Determining the PerR regulon allowed to identify the PerR-dependent mechanisms of the peroxide adaptive response and has revealed a PerR-independent regulatory network involving other transcriptional regulators, two-component systems and sigma factors as well as non-coding RNAs that putatively orchestrate, in concert with PerR, this adaptive response. In addition, we have identified other PerR-regulated genes encoding a TonB-dependent transport system, a lipoprotein (LipL48) and a two-component system (VicKR) involved in *Leptospira* tolerance to superoxide and that could represent the first defense mechanism against superoxide in *L. interrogans*, a bacterium lacking canonical superoxide dismutase. Our findings provide a comprehensive insight into the mechanisms required by pathogenic *Leptospira* to overcome infection-related oxidants during the arm race with a host. This will participate in framing future hypothesis-driven studies to identify and decipher novel virulence mechanisms in this life-threatening pathogen.

**Author summary:** Leptospirosis is a zoonotic infectious disease responsible for over one million of severe cases and 60 000 fatalities annually worldwide. This neglected and emerging disease has a worldwide distribution, but it mostly affects populations from developing countries in sub-tropical areas. The causative agents of leptospirosis are pathogenic bacterial *Leptospira* spp. There is a considerable deficit in our knowledge of these atypical bacteria, including their virulence mechanisms. During infection, *Leptospira* are confronted with the deadly oxidants produced by the host tissues and immune response. Here, we have identified the cellular factors necessary for *Leptospira* to overcome the oxidative stress response. We found that *Leptospira* solicit peroxidases to detoxify oxidants as well as chaperones of the heat shock response and DNA repair proteins of the SOS response to recover from oxidative damage. Moreover, our study indicates that adaptation to oxidative stress is orchestrated by a regulatory network involving PerR and other transcriptional regulators, sigma factors, two component systems, and putative non-coding RNAs. These findings provide a comprehensive insight into the mechanisms required by pathogenic *Leptospira* to tolerate infection-related oxidants, helping identify novel virulence factors, developing new therapeutic targets and vaccines against leptospirosis.

## Introduction

In order to invade a host and establish persistent colonization, pathogens have evolved a variety of strategies to resist, circumvent, or counteract host defenses. Synthesis of detoxification enzymes or molecules to eliminate host-produced bactericidal compounds, secretion of effectors inhibiting or subverting the host innate immunity, biofilm formation enabling resistance to host defenses, are all examples of mechanisms used by pathogens depending of their lifestyle and niche.

The whole strategies used by pathogenic *Leptospira* for successful host colonization and virulence are not fully unraveled. These aerobic gram-negative bacteria of the spirochetal phylum are the causative agents of leptospirosis, a widespread zoonosis (1). Although recognized as a health threat among impoverished populations in developing countries and tropical areas (2), reported cases of leptospirosis are also on the rise in developed countries under temperate climates (3). Rodents are the main reservoir for leptospires as the bacteria asymptomatically colonize the proximal renal tubules of these mammals. They shed bacteria in the environment by their urine and leptospires are transmitted to other animals and humans mostly by exposure to contaminated soils and water. Once having penetrated an organism, *Leptospira* enter the bloodstream and rapidly disseminate to multiple tissues and organs including kidney, liver and lungs. Clinical manifestations range from a mild flu-like febrile state to more severe and fatal cases leading to hemorrhages and multiple organ failure. The lack of efficient tools and techniques for genetic manipulation of *Leptospira* spp. and their fastidious growth in laboratory conditions have greatly hampered and limited our understanding of their mechanisms of pathogenicity and virulence (4,5).

As part of the host innate immunity response, reactive oxygen species (ROS), *i.e.* superoxide anion (·O_2_^-^), hydrogen peroxide, (H_2_O_2_), hydroxyl radicals (·OH), hypochlorous acid (HOCl), and nitric oxide anion (·NO) are produced upon infection by *Leptospira*. Indeed, the internalization of pathogenic *Leptospira* by macrophages and concomitant production of these oxidants have been demonstrated *in vitro* (6), and leptospirosis-associated oxidative stress has been observed in leptospirosis patients (7) and infected animals (8). Consistent with these findings was the demonstration that catalase, that catalyzes the degradation of H_2_O_2_, is required for *Leptospira interrogans* virulence (9).

Pathogenic *Leptospira* spp. are among the rare examples of gram-negative bacteria where defenses against peroxide stress, such as catalase, are controlled by a peroxide stress regulator (PerR) and not by OxyR (10). PerR is a peroxide-sensing transcriptional repressor that belongs to the Fur (Ferric uptake regulator) family of regulators, mostly present in gram-positive bacteria (11). The *B. subtilis* PerR is in a DNA-binding prone conformation in the presence of a regulatory metal (Fe^2+^) (12). Upon oxidation by H_2_O_2_, PerR releases its regulatory metal and switches to a conformation that cannot bind DNA, leading to the alleviation of gene repression (13,14).

We have conducted a structural and functional characterization of PerR in *L. interrogans* and showed that *Leptospira* PerR exhibits the typical metal-induced conformational switch controlling DNA binding and release (15). Our findings indicated that not only *Leptospira* PerR represses defenses against H_2_O_2_, but also a *perR* mutant had a decreased fitness in other host-related stress conditions including in the presence of superoxide (15). Interestingly, it was shown that *perR* is up-regulated when *Leptospira* are exposed *in vitro* to hydrogen peroxide (15) as well as when *Leptospira* are cultivated *in vivo* using Dialysis Membrane Chambers (DMCs) in rats (16), which strongly suggests a role of PerR in the adaptation of pathogenic *Leptospira* to a mammalian host.

In order to identify the mechanisms solicited by pathogenic *Leptospira* to adapt to oxidative stress, we have determined the global transcriptional response of *L. interrogans* to H_2_O_2_ and assessed the role of PerR in this adaptation. This has revealed the cellular factors constituting the first-line of defense against ROS that *Leptospira* might encounter when infecting a mammalian host. In addition, our study has identified repair mechanisms allowing leptospires to recover from oxidative damage. Putative regulatory non-coding RNAs were also pinpointed, indicating the complexity of the regulatory network controlling the adaptive response to peroxide. We have also identified novel PerR-regulated factors involved in *Leptospira* survival in the presence of superoxide and assessed their role in *Leptospira* virulence.

## Results

### *Leptospira* transcriptional response to a sublethal concentration of hydrogen peroxide

In order to characterize the transcriptional response of pathogenic *Leptospira* to hydrogen peroxide, we have exposed exponentially growing *L. interrogans* cells to sublethal concentrations of this oxidant. A 30 min treatment with 10 µM H_2_O_2_ (in the presence of iron) was chosen during pilot experiments as having no significant effect on *Leptospira* viability and growth during logarithmic phase while increasing expression of H_2_O_2_-responsive genes such as *perR* (15). RNA sequencing was performed to assess RNA abundance and comparison with untreated cells identified a total of 21 genes with differential transcript abundance. Among those, only 12 and 1 genes were respectively up- and down-regulated by a at least two-fold with P-values ≤0.005 (See Table 1).

**Table 1:** Differentially expressed genes upon exposure to sublethal dose of H_2_O_2_.

| ORF ID<sup>a</sup> | Gene | Function | Log<sub>2</sub>FC | Adjusted p-value | FC (RT-qPCR)<sup>b</sup> |
| --- | --- | --- | --- | --- | --- |
| **Up-regulated genes** |  |  |  |  |  |
| LIMLP_02795 (LIC12927/LA0666) | *ccp* | Cytochrome C peroxidase | 4.764* | 5.31e-43 | 38.900 |
| LIMLP_05955 (LIC11219/LA2809) | *ahpC* | Peroxiredoxin/alkylperoxiredoxin reductase | 3.145* | 3.63e-20 | 11.742 |
| LIMLP_05960 (LIC11220/LA2808) | *sufB* | Fe-S cluster assembly protein | 1.056* | 1.60e-08 | 1.880 |
| LIMLP_10145 (LIC12032/LA1859) | *katE* | Catalase | 1.786 | 2.11e-08 | 3.477 |
| LIMLP_10150 (LIC12033/LA1858) |  | Ankyrin repeat-containing protein | 2.051* | 2.30e-11 | 4.183 |
| LIMLP_10155 (LIC12034/LA1857) | *perR* | Regulator Fur family | 2.319* | 1.02e-39 | 6.827 |
| LIMLP_17840 (LIC20008/LB010) | *hemA* | Glutamyl-tRNA reductase | 1.771* | 1.16e-10 | 3.389 |
| LIMLP_17845 (LIC20009/ LB011) | *hemC/D* | Porphobilinogen deaminase | 1.617* | 6.57e-13 | 2.328 |
| LIMLP_17850 (LIC20010/ LB012) | *hemB* | Delta-aminolevulinic acid dehydratase | 1.455* | 2.65e-14 | 2.064 |
| LIMLP_17855 (LIC20011/ LB013) | *hemL* | Glutamate-1 semialdehyde aminotransferase | 1.262* | 1.19e-07 | 2.193 |
| LIMLP_17860 (LIC20012/ LB014) |  | Signal transduction histidine kinase | 1.035* | 2.67e-03 | 2.470 |
| LIMLP_17865 (LIC20013/ LB015) |  | Response regulator cheY | 1.166* | 1.01e-03 | 2.012 |
| **Down-regulated genes** |  |  |  |  |  |
| LIMLP_18600 (LIC20149/ LB187) |  | Permease of the Major facilitator superfamily | -1.001 | 1.17e-04 | 0.894 |

Significantly up-and down-regulated genes upon a 30 min exposure to 10 μM H<sub>2</sub>O<sub>2</sub> with a Log<sub>2</sub>FC cutoff of 1 and an adjusted p-value cutoff of 0.005.
<sup>a</sup> Gene numeration is according to Satou *et al.* (52). Corresponding genes of *L. interrogans* serovar lai strain 56601 and serovar Copenhageni Fiocruz strain L1-130 are indicated in parenthesis.
<sup>b</sup> Fold change in gene expression upon a 30 min exposure to 10 μM H<sub>2</sub>O<sub>2</sub> obtained by RT-qPCR experiments.
\* Genes significantly up-and down-regulated by Volcano analysis (Log<sub>2</sub>FC cutoff of 1 and p-value cutoff of 0.005 as seen in Fig 1A).

Under a low concentration of H_2_O_2_, LIMLP_10145, encoding a catalase, and LIMLP_02795 and LIMLP_05955, coding respectively for a cytochrome C peroxidase and for a peroxiredoxin, were up-regulated with a Log_2_FC of 1.79, 4.76 and 3.14, respectively.

The catalase encoded by LIMLP_10145 (*katE*) is a monofunctional heme-containing hydroperoxidase, the catalase activity of which and periplasmic localization were experimentally demonstrated in pathogenic *Leptospira* (9,17,18). The immediate upstream ORF (LIMLP_10150), encoding an ankyrin repeat-containing protein, was also up-regulated with a comparable fold. In bacteria such as *Pseudomonas aeruginosa* and *Campylobacter jejuni*, protein with ankyrin repeats were found to be required for the catalase activity, probably by allowing heme binding (19,20). In *L. interrogans, katE* and *ank* were organized as an operon (S1 Fig) and significant up-regulation of the *ank-katE* operon upon exposure to sublethal dose of H_2_O_2_ was confirmed by RT-qPCR (Table 1 and S1 Fig).

The significantly up-regulated *ahpC* gene (LIMLP_05955) encodes a peroxiredoxin that reduces H_2_O_2_ and ter-butyl peroxide (21). The SufB-encoding LIMLP_05960 located in the vicinity of *ahpC* was also up-regulated with a 2-fold. *SufB* encodes a polypeptide involved in Fe-S cluster assembly proteins. In bacteria such as *Escherichia coli*, SufB is part of a complex composed of SufB, SufD and the SufC ATPase. *SufB* is normally found in an operon with *sufC* and *sufD* as well as with the other factors of the Suf machinery, i. e. *sufE* and the *sufS* (encoding cysteine desulfurases). *L. interrogans* genome contains a putative *suf* cluster (LIMLP_14560-14580 ORFs) and SufE (encoded by LIMLP_05090) but none of the other *suf* ORFs were regulated by sublethal dose of H_2_O_2_. The SufB-encoding LIMLP_05960 shares 40% and 47% identity with SufB from *E. coli* and *B. subtilis*, respectively, and most importantly it does contain the critical Cysteine residue suggesting that the isolated LIMLP_05960-encoded SufB functions as a genuine scaffold in Fe-S biogenesis. A function or cooperation of SufB with AhpC in H_2_O_2_ detoxification remains to be demonstrated.

LIMLP_02795 was another peroxidase-encoding ORF that was greatly up-regulated in the presence of H_2_O_2_ (Log_2_FC of 4.76). LIMLP_02795 encodes a putative Cytochrome C Peroxidase (CCP) family that catalyzes the reduction of H_2_O_2_ into H_2_O using the ferrocytochrome as an electron donor. In several *L. interrogans* genomes, this ORF is annotated as a MauG, a class of Cytochrome C Peroxidase that catalyzes the oxidation of methylamine dehydrogenase (MADH) into tryptophan tryptophylquinone (TTQ) in the methylamine metabolism pathway. LIMLP_02795 exhibits two heme domains with the conventional heme binding motif CXXCH that exists in both CCP and MauG proteins, but it lacks the Tyrosine axial ligand for heme (Tyr294 in *Paracoccus denitrificans*, (22)) that is conserved in all MauGs and replaced by a Methionine or Histidine residue in CCPs. Therefore, it is very likely that LIMLP_02795 encodes a CCP with a peroxidase activity that is not involved in the methylamine metabolism pathway.

In addition to these three peroxidases, whose increased expression was confirmed by RT-qPCR (Table 1 and S1 and S2 Figs), several ORFs encoding components of heme biosynthesis (LIMLP_17840-17865) were up-regulated by a 2 to 3.4-fold (Table 1). *Leptospira*, unlike other spirochetes, possess a complete heme biosynthesis functional pathway (23). The ORFs encoding the glutamyl-tRNA reductase (*hemA*), porphobilinogen deaminase (*hemC/D*), delta-aminolevulinic acid dehydratase (*hemB*), glutamate-1 semialdehyde aminotransferase (*hemL*), uroporphyrinogen-III decarboxylase (*hemE*), coproporphyrinogen-III oxidase (*hemN/F*), as well a two-component system (TCS) (LIMLP_17860 and LIMLP_17865) were organized as an operon (S3 Fig). RT-qPCR confirms the significance of the up-regulation of *hemA, hemC/D* and of the LIMLP_17860-encoded histidine kinase of the TCS (Table 1 and S3 Fig).

When pathogenic *Leptospira* cells are exposed to 10 µM H_2_O_2_, the only ORF that was down-regulated was that encoding a permease (LIMLP_18600, with a Log_2_FC of −1). This permease is a putative Major Facilitator Superfamily (MFS) transporter and is predicted to contain 12 transmembrane helixes. This permease-encoding ORF is the second gene of a bicistronic operon where a heme oxygenase-encoding ORF (LIMLP_18595) is the first (S4 Fig). Expression of the heme oxygenase ORF was not significantly changed by the exposure to 10 µM H_2_O_2_ (S4 Fig).

Plotting statistical significance (-log10 of p-values) in function of fold change (Log_2_FC) confirmed that *katE, ccp, ahpC, perR*, and several genes of the heme biosynthesis pathway were among the genes the expression of which was significantly up-regulated (Fig 1A).

**Fig 1.**
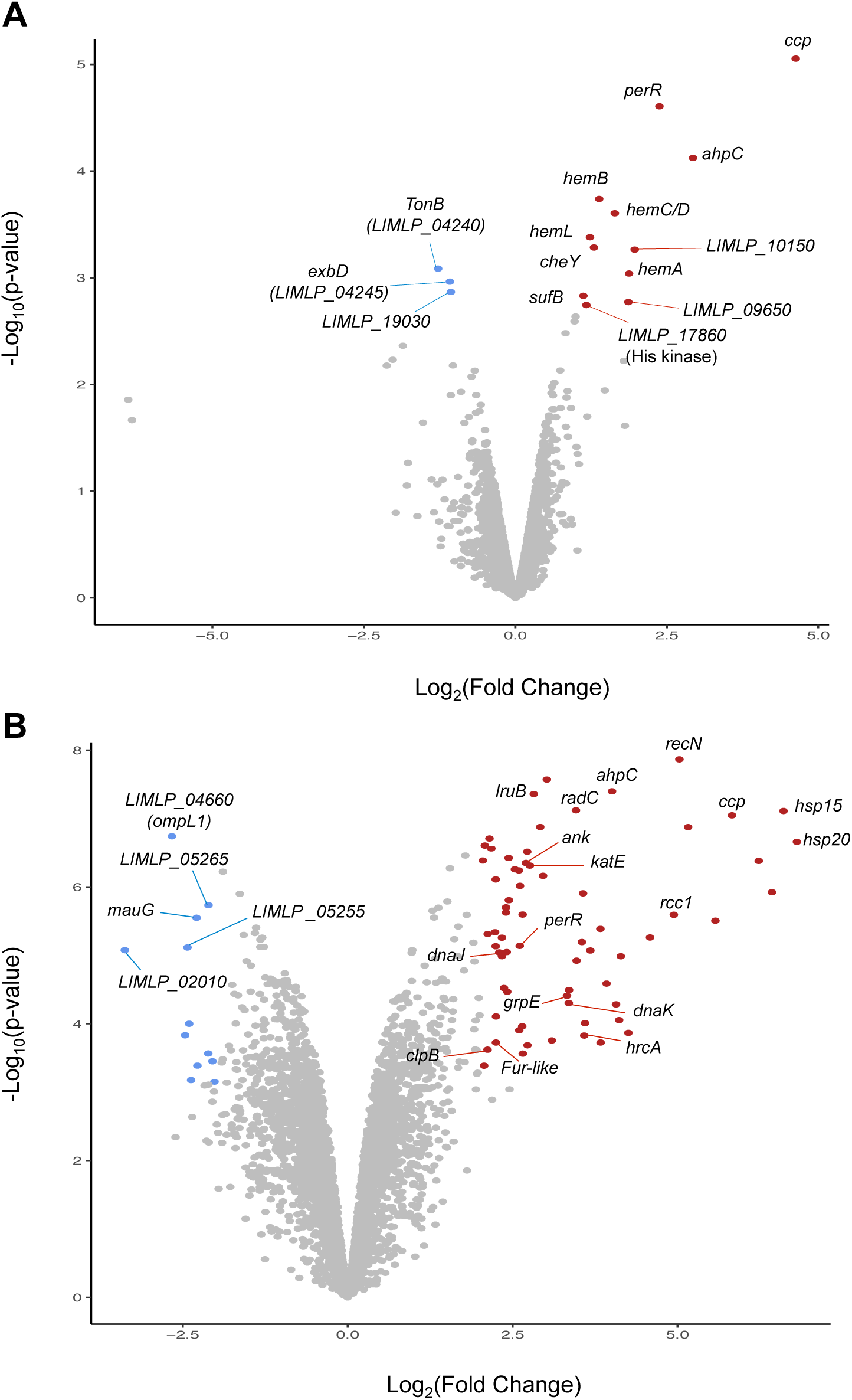
Volcano representation of differentially expressed genes upon exposure to hydrogen peroxide. Up- and down-regulated genes upon a 30 min exposure to 10 µM H_2_O_2_ (A) or 1-hour exposure to 1 mM H_2_O_2_ (B) were graphically represented by a Volcano analysis. Red and blue dots indicate up- and down-regulated genes, respectively, with significant change in expression with a Log_2_FC cutoff of 1 and p-value<0.05 in (A) and with a Log_2_FC cut off of 2 and p-value<0.05 in (B). A higher Log_2_FC cutoff were chosen for the data obtained upon exposure to 1 mM H_2_O_2_ (in B) due to a much greater number of differentially expressed genes. Representative genes are labeled.

Noteworthily, after a 2-hour exposure of *L. interrogans* to 10 µM H_2_O_2_, the expression of the peroxidases and heme biosynthesis genes returns to a level closer to that observed in the absence of H_2_O_2_ (S1-S4 Figs). Altogether, these data indicate that pathogenic *Leptospira* respond to a sublethal dose of H_2_O_2_ by soliciting three peroxidases and heme, and that the up-regulated peroxidase and catalase activities are sufficient to allow survival of *Leptospira*.

### *Leptospira* transcriptional response to 1 mM of hydrogen peroxide

In order to better reproduce harmful oxidative stress encountered during infection, we performed similar RNASeq experiments upon 1-hour exposure to 1 mM H_2_O_2_. In this condition, *Leptospira* survival was of 60% ± 2.735 as assessed by plating on EMJH agar plates. Comparison with untreated cells identified a total of 2145 genes with differential transcript abundance. Among those, 223 and 268 genes were respectively up- and down-regulated by a ≥2.0 fold with p-values ≤0.0005. The volcano representation exhibited more scattered data points (Fig 1B), bearing witness to a higher number of genes with significantly and statistically changed expression than when *Leptospira* are exposed to sublethal dose of H_2_O_2_ (Fig 1A).

Differentially expressed genes were classified into COG functional categories and the obtained COG frequencies were compared to the frequency of the genes in the genome. As seen in Fig 2, the up-regulated genes were enriched in the post-translational modification, protein turnover, and chaperones categories whereas down-regulated genes mainly fell into metabolism, translation and ribosomal structure and biogenesis, coenzyme transport and metabolism, and energy production and conversion categories.

**Fig 2.**
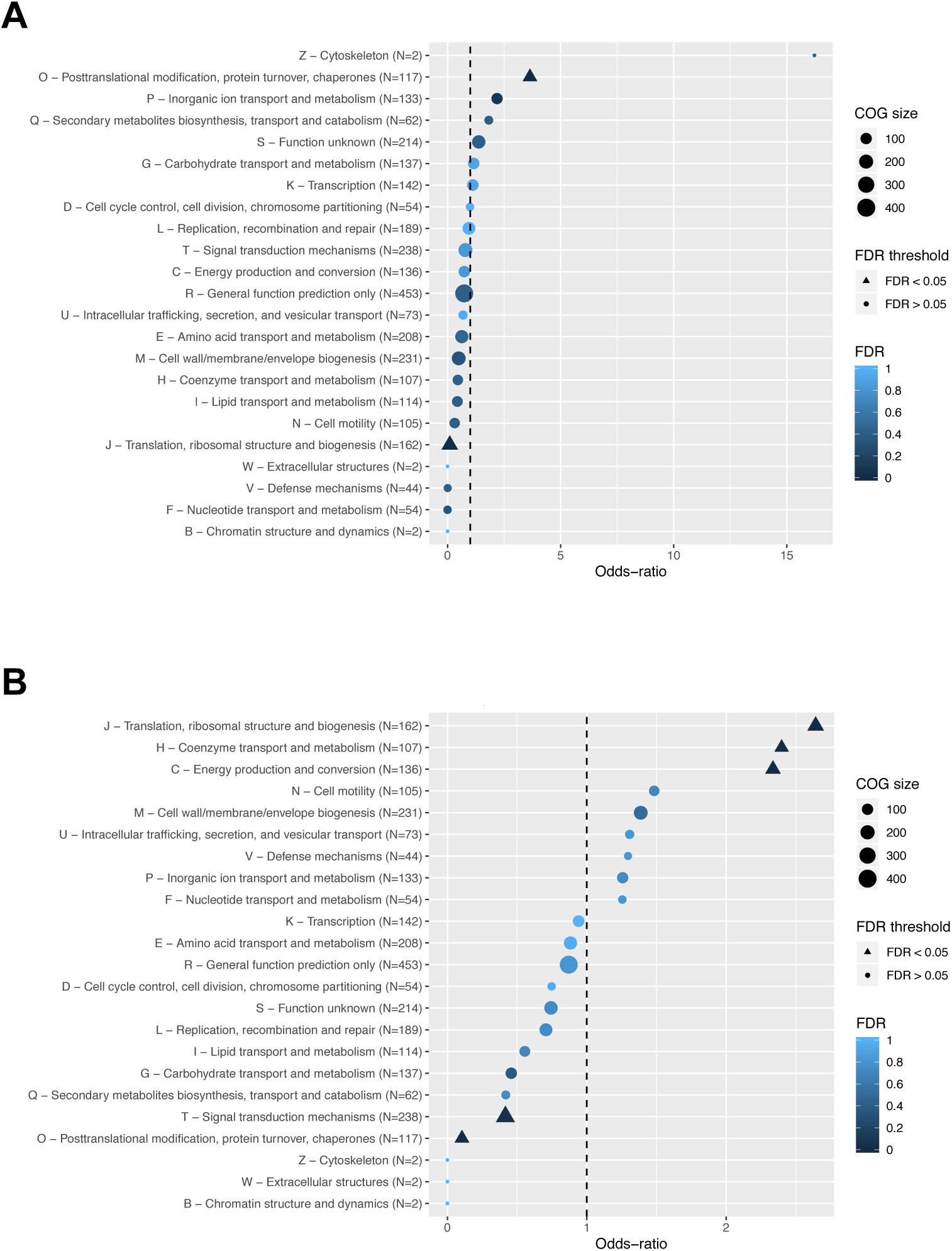
Classification of differentially expressed genes upon exposure to lethal dose of hydrogen peroxide. ORFs with significantly changed expression when *L. interrogans* were exposed to 1 mM H_2_O_2_ for 1h were classified according to the COG (Clusters of Orthologous Groups). A Log_2_FC cutoff of 1 and −1 were applied for the up-regulated (A) and down-regulated (B) ORFs, respectively, with an adjusted p-value<0.005. An odd-ratio higher or lower than 1 indicates an over- or under-representation of a functional category, respectively, and a COG category with a False Discovery Rate (FDR) lower than 5 % is considered as enriched. The functional categories are indicated on the left.

As in the presence of low dose of H_2_O_2_, the *ank-katE* operon (LIMLP_10150-10145), *ccp* (LIMLP_02790) and *ahpC* (LIMLP_05955) were up-regulated in the presence of 1 mM H_2_O_2_ but with higher fold changes (with Log_2_FC values of 2.7, 5.8 and 4, respectively, see Fig 3 and S1 Table). Noteworthy, the ORF upstream *ahpC* that encodes a SufB (LIMLP_05960) was also significantly up-regulated (with a with Log_2_FC value of 2.2). Likewise, *perR* expression was greater in the presence of 1 mM H_2_O_2_ (with Log_2_FC value of 3.5, see Fig 3 and S1 Table). All these up-regulations were confirmed by RT-qPCR experiments (S1 Table). Additional ORFs encoding cellular factors related to oxidative stress and redox maintenance were also up-regulated (Fig 3 and S1 Tables). An ORF encoding a thiol oxidoreductase (LIMLP_07145) exhibiting two cytochrome C-like (heme binding) domains was up-regulated with a Log_2_FC value of 2.2. LIMLP_07145 was located immediately downstream an ORF (LIMLP_07150) encoding a protein with five chromosome condensation regulator (RCC1) domains that was up-regulated with a Log_2_FC value of about 5. LIMLP_07145-07150 are probably organized as a bicistronic operon as predicted in Zhukova *et al.* (24). A second thiol peroxidase-encoding ORF (LIMLP_14175) exhibiting a single cytochrome C-like domain was also up-regulated (Log_2_FC value of 1.8). This ORF might be part of the operon LIMLP_14170-14180 where LIMLP_14170 and LIMP_14180, two ORFs annotated as Imelysins (iron-regulated proteins), were also up-regulated (Log_2_FC value of 2.8 and 1.4, respectively). Noteworthily, the Imelysin encoded by LIMLP_14170 is the LruB protein that was shown to be associated with *Leptospira*-induced uveitis (25).

**Fig 3.**
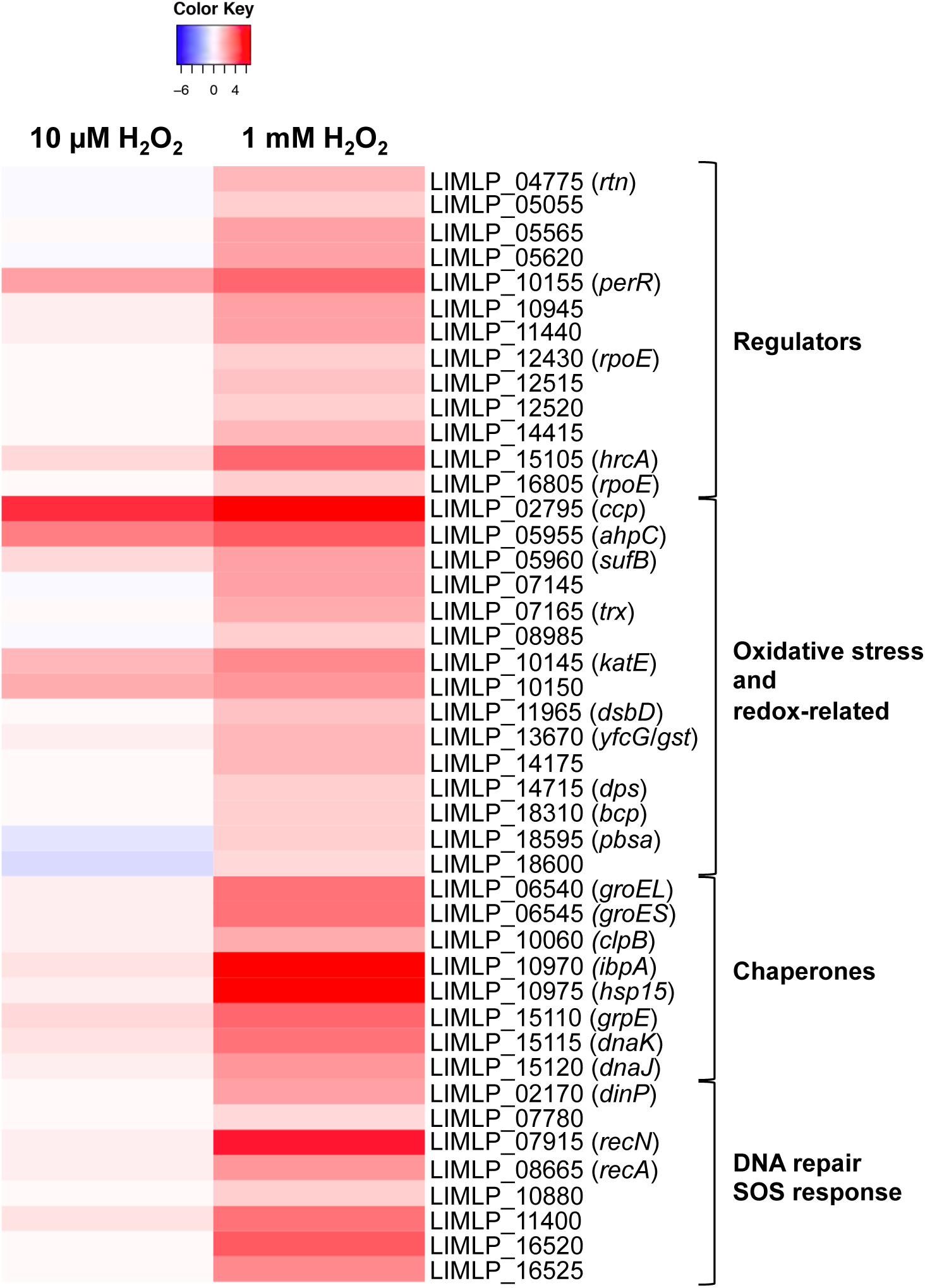
Comparison of up-regulated genes upon exposure to sublethal and lethal dose of hydrogen peroxide. The expression of selected up-regulated genes determined by RNASeq when *L. interrogans* are exposed 1 hour to 1 mM H_2_O_2_ was compared to that of *L. interrogans* exposed 30 min to 10 µM H_2_O_2_. Genes are organized by their function, and their number and name are indicated on the right. The Heat Map color from blue to red indicates low to high Log_2_FC (with H_2_O_2_ versus without H_2_O_2_).

A thioredoxine disulfide reductase (encoded by LIMLP_07165) was up-regulated (Log_2_FC value of 1.9, see Fig 3 and S1 Table). This protein has been shown to catalyze *in vitro* the NADPH-dependent reduction of a thioredoxin encoded by LIMLP_09870 (26). The LIMLP_09870 was only slightly up-regulated in the presence of 1 mM H_2_O_2_ (Log_2_FC value of 0.8, see Table S2).

Other thiol peroxidase-encoding ORFs were up-regulated, including LIMLP_08985 that encode a glutaredoxin, LIMLP_11965 that codes for the thiol disulfide interchange protein DsbD that might participate in the oxidative folding of periplasmic proteins, and LIMLP_18310 that encodes a bacterioferritin comigratory protein (Bcp) (See Fig 3 and S1 Table). An ORF encoding a putative Glutathione S transferase (LIMLP_13670) had an increased expression in the presence of 1 mM H_2_O_2_ (Log_2_FC value of 1.76), as did an ORF annotated as DNA binding stress protein (Dps) (Log_2_FC value of 1.09) (See Fig 3 and S1 Table).

Major cellular pathways involved in reparation of damaged cellular components were dramatically up-regulated when *Leptospira* were exposed to a lethal dose of H_2_O_2_. Indeed, several genes encoding molecular chaperones had an increased expression in the presence of 1 mM H_2_O_2_ (Fig 3 and S1 Table). Two ORFs encoding small heat shock proteins (sHSP), probably organized as a bicistronic operon (LIMLP_10970-10975), exhibited a significant increase in expression (Log_2_FC values of about 6). The LIMLP_15105-15120 cluster encoding the DnaK/DnaJ/GrpE molecular chaperone machinery and its putative repressor HrcA, was significantly up-regulated with Log_2_FC values of 2.6-3.6. Similarly, the GroES-GroEL operon (encoded by LIMLP_06545-06540) was up-regulated with a Log_2_FC values of 3.3. The *clpB* gene (LIMLP_10060) also had an increased expression (Log_2_FC value of 2.1). Thus, the machinery necessary for preventing protein aggregation and promoting protein refolding is solicited when *Leptospira* are exposed to high dose of H_2_O_2_.

Genes encoding several components of the SOS response, a regulatory network stimulated by DNA damage-inducing stress, had a higher expression in the presence of 1 mM H_2_O_2_ (Fig 3 and S1 Table). Indeed, ORFs encoding the recombinase A (*recA*, LIMLP_08665), the DNA repair protein RecN (LIMLP_07915), the DNA polymerase IV (*dinP*, LIMLP_02170) as well as the repressor of the SOS response LexA1 (LIMLP_11440) were significantly up-regulated. Other factors putatively involved in DNA repair but not under the control of LexA1 (27,28) had also an increased expression, including the DNA mismatch repair protein MutS (LIMLP_07780, Log_2_FC value of 1) and the DNA repair protein RadC (LIMLP_11400, Log_2_FC value of 3.4).

One remarkably up-regulated ORF (LIMLP_00895) was located into a genomic region previously identified as an island enriched in prophage genes ranging from LIMLP_00855 to LIMLP_01005 and referred as prophage 1 (28,29) (S1 Table). Also, another cluster enriched in prophage genes (from LIMLP_13010 to LIMLP_13095), referred as prophage 2 (28), contains 4 ORFs (LIMLP_13010, LIMLP_13015, LIMLP_13020, and LIMLP_13025) that were up-regulated in the presence of 1 mM H_2_O_2_ (S1 Table).

Down-regulated genes were mainly genes putatively involved in translation and metabolism (Fig 2B and S2 Table). 14 ORFs encoding ribosomal proteins, a translation initiation factor (LIMLP_03190), a ribosome maturation factor (LIMLP_07600), a RNA polymerase RpoA (LIMLP_03215), a transcription termination factor RhoA (LIMLP_13190) were among them. A cluster of gene encoding the ATP synthase complex (LIMLP_06050-06080) was down regulated with a Log_2_FC≤-1.2, indicating that *Leptospira* decrease ATP synthesis upon exposure to high dose of H_2_O_2_ (S2 Table). Another metabolic pathway that was down-regulated in this condition was the cobalamin (vitamine B12) biosynthesis pathway. Indeed, 15 out 17 genes of the cobI/III cluster (LIMLP_18460-18530) were significantly down-regulated (with a Log_2_FC ≤-1.5, S2 Table).

A cluster of four genes encoding proteins of the CRISPR-CAS machinery (*csh2*, LIMLP_2870; *cas8*, LIMLP_2875; *cas5*, LIMLP_2880; *cas3*, LIMLP_2880) putatively involved in phage defense were down-regulated (with a Log_2_FC<1, see S2 Table).

Finally, several genes related to motility/chemotaxis were down-regulated when *Leptospira* are exposed to a high dose of H_2_O_2_. Several of these genes encode constituents of the endoflagellum basal body (*flgGAHIJ*, LIMLP_06485-06505), of the flagellar export apparatus (*fliOPQR-FlhBA*, LIMLP_06690-06715; *fliL*, LIMLP_14615 and LIMLP_14620), and of the flagellar motor stator (*motAB*, LIMLP14625-14630), and chemotaxis-related proteins (*cheBDW*-mcp, LIMLP_07420-07435).

### Identification of differentially non-coding RNAs in the presence of hydrogen peroxide

In order to identify non-coding RNAs (ncRNAs) whose expression is changed in the presence of hydrogen peroxide, non-coding genome regions of RNASeq data were also analyzed. When *Leptospira* were exposed to 10 *μ*M H_2_O_2_ for 30 min, only a few ncRNAs were differentially expressed. The most highly up-regulated ncRNA was a 322 bp ncRNA (rh859, Log_2_FC of 3.19) located 21 bp downstream LIMLP_02795, the ORF encoding the cytochrome C peroxidase. Other significantly up-regulated ncRNAs were a 127 bp ncRNA (rh3130, Log_2_FC of 1.92) located downstream the two small Hsp-encoding ORFs (LIMLP_10970-10975) and a 225 bp ncRNA (rh3999, Log_2_FC of 1.90) overlapping with the LIMLP_14135 ORF.

When *Leptospira* were exposed to a lethal dose of hydrogene peroxide (1 mM for 1h), a higher number of differentially expressed ncRNAs was detected. Indeed, 416 and 102 ncRNAs were up- and down-regulated, respectively. 28 ncRNAs were up-regulated with a Log_2_FC above 1.5. Rh3130 and rh3352 were the two most highly up-regulated ncRNAs with a Log_2_FC > 7 (Table 2). A 70 bp ncRNA (rh288), which was up-regulated with a Log_2_FC of 3.81, overlapped with LIMLP_00895, an ORF located in the prophage locus 1 (Table 2). 13 ncRNAs were down-regulated with a Log_2_FC below −1.5. Among the most highly down-regulated ncRNAs was a 193 bp RNA (rh967, Log_2_FC of −2.66) located downstream a large cluster encoding ribosomal proteins (LIMLP_03075-3220).

**Table 2:**
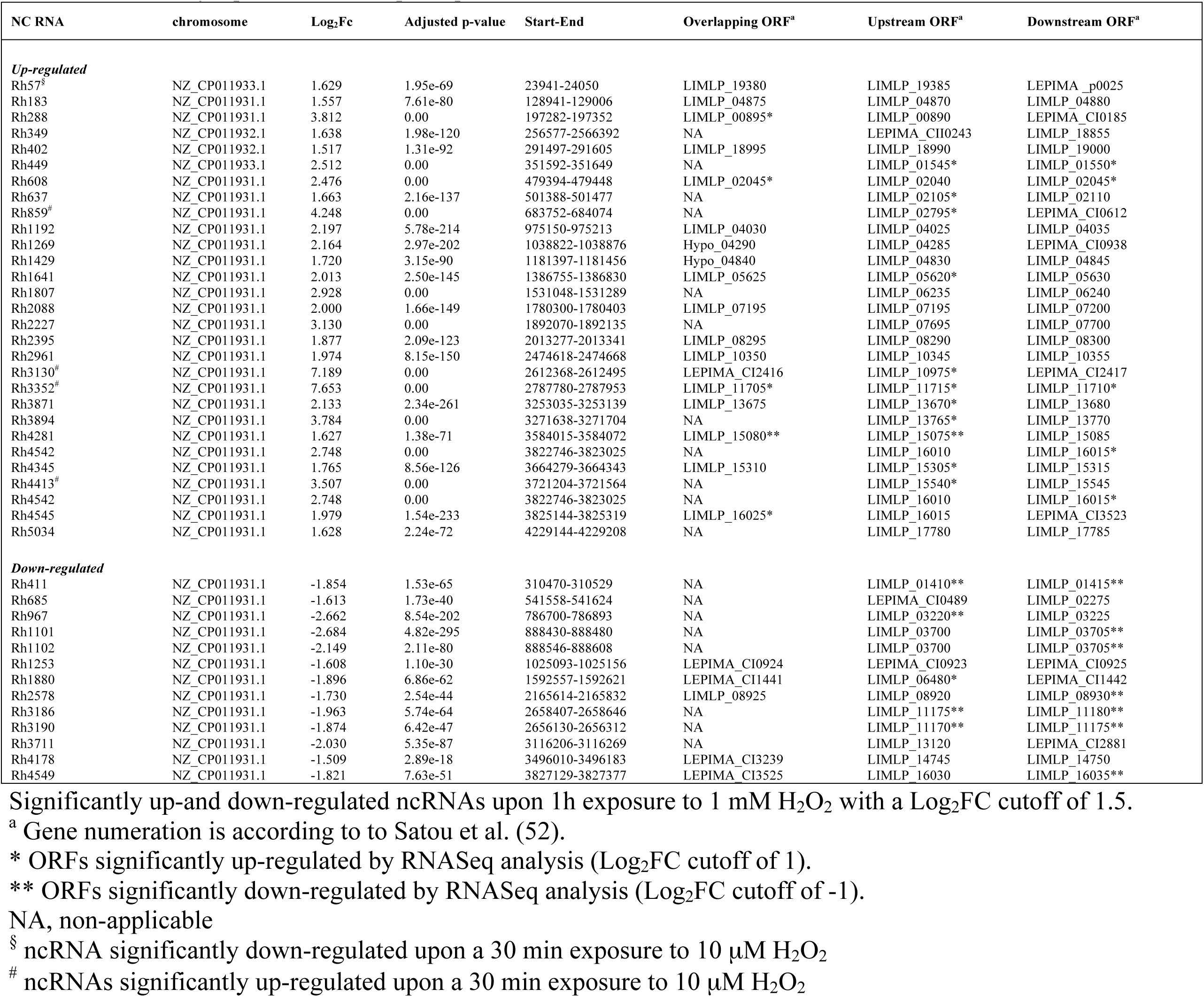
Differentially expressed ncRNAs upon exposure to lethal dose of H_2_O_2_.

Several of the ncRNAS whose expression was up- or down-regulated in the presence of hydrogen peroxide were located in the vicinity or overlapped ORFs that were also up- or down-regulated in the same conditions. For instance, the rh3130 and rh859, two of the most highly up-regulated ncRNAs, were in the vicinity of Hsp20 and CCP-encoding ORFs (LIMLP_10970-10975 and LIMLP_02795, respectively), two genes whose expression was greatly increased in the presence of hydrogen peroxide (Tables 1 and 2 and Fig 3). LIMLP_05620, LIMLP_13670, and LIMLP_13765 were three up-regulated ORFs upon exposure to hydrogen peroxide that have a up-regulated downstream ncRNA (rh1641, rh3871, and rh3894, respectively). This tendency was also observed with down-regulated ncRNAs. Rh411, rh967, rh1101, rh1102, rh1880, rh3186, and rh4281 ncRNAs were also located downstream or upstream, or overlapped ORFs whose expression was decreased in the presence of hydrogen peroxide (Table 2).

Most of the predicted ncRNAs show little homology with well characterized RNAs families of the RFam database. However, this study has allowed the identification of a TPP riboswitch (rh1210, RFam 00023) in the intergenic region of LIMLP_04090 and LIMLP_04095 (rh1210) and a cobalamin riboswitch (RFam 00174) in the intergenic regions of LIMLP_06575 and LIMLP_06580 (rh1913), of LIMLP_11825 and LIMLP_11830 (rh3382), and upstream LIMLP_17140 (rh4854). An AsrC (Antisense RNA of rseC, RFam 02746) was overlapping with LIMLP_10025 (rh2876) and a ligA thermometer (RFam02815) was located in the intergenic region of LIMLP_05075 and LIMLP_05080 (rh1488). None of those ncRNAs were significantly differentially expressed upon *Leptospira* exposure to H_2_O_2_.

Altogether, these findings indicate that exposure of *Leptospira* to 1 mM H_2_O_2_ triggers a drastic regulation of ncRNA expression that correlates with dramatic changes in coding sequence expression.

### Contribution of PerR in the adaptation of pathogenic *Leptospira* to oxidative stress

Comparison of the transcriptome of a *perR* mutant with that of WT strain allowed determination of the PerR regulon in *L. interrogans*. In the *perR* mutant, 5 and 13 ORFs were up- and down-regulated, respectively, with a log_2_FC cutoff of 1 and a p-value below 0.05 (Table 3).

**Table 3:**
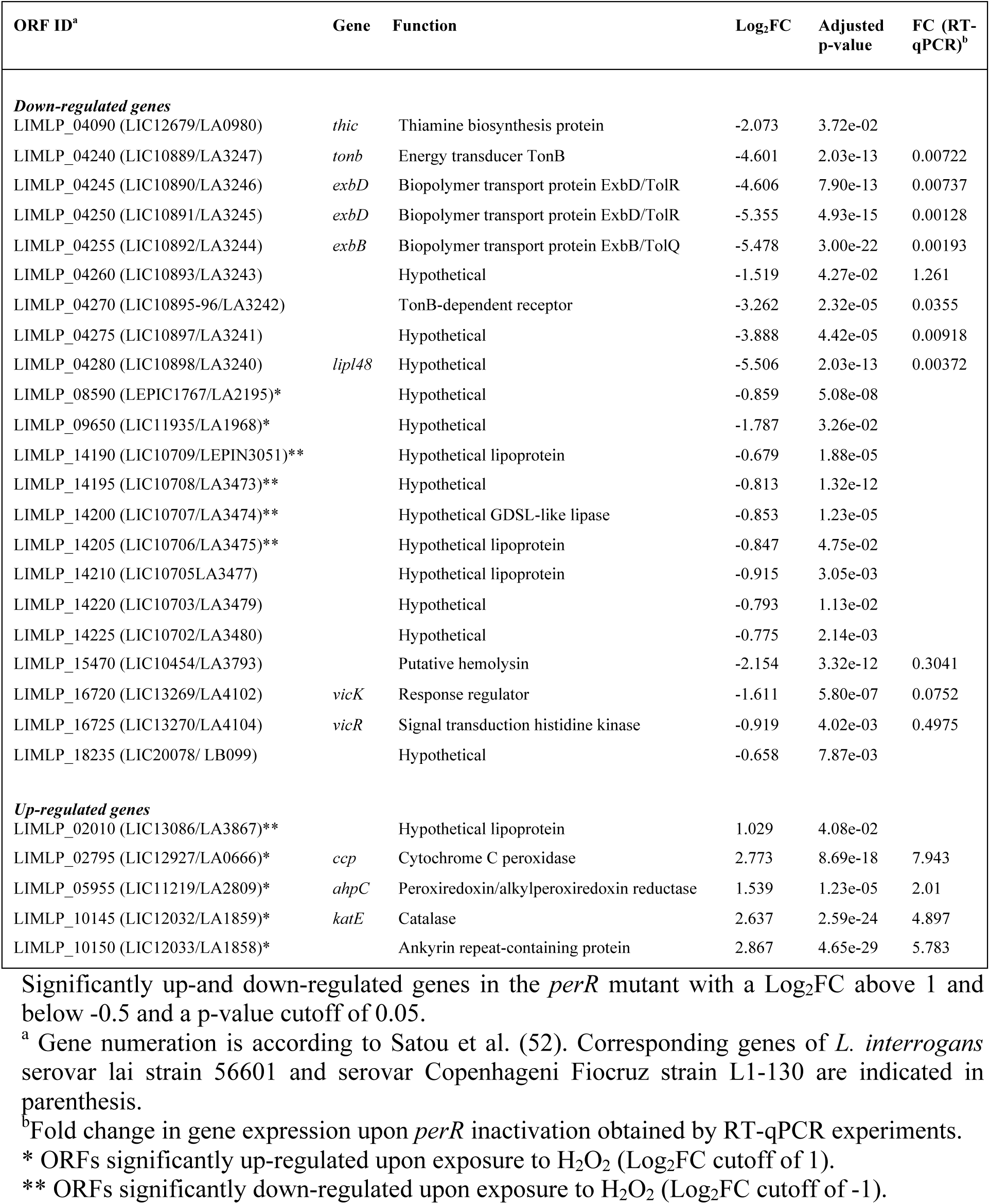
Differentially expressed genes upon *perR* inactivation.

| ORF ID<sup>a</sup> | Gene | Function | Log<sub>2</sub>FC | Adjusted p-value | FC (RT-qPCR)<sup>b</sup> |
| --- | --- | --- | --- | --- | --- |
| **Down-regulated genes** |  |  |  |  |  |
| LIMLP_04090 (LIC12679/LA0980) | *thic* | Thiamine biosynthesis protein | -2.073 | 3.72e-02 |  |
| LIMLP_04240 (LIC10889/LA3247) | *tonb* | Energy transducer TonB | -4.601 | 2.03e-13 | 0.00722 |
| LIMLP_04245 (LIC10890/LA3246) | *exbD* | Biopolymer transport protein ExbD/TolR | -4.606 | 7.90e-13 | 0.00737 |
| LIMLP_04250 (LIC10891/LA3245) | *exbD* | Biopolymer transport protein ExbD/TolR | -5.355 | 4.93e-15 | 0.00128 |
| LIMLP_04255 (LIC10892/LA3244) | *exbB* | Biopolymer transport protein ExbB/TolQ | -5.478 | 3.00e-22 | 0.00193 |
| LIMLP_04260 (LIC10893/LA3243) |  | Hypothetical | -1.519 | 4.27e-02 | 1.261 |
| LIMLP_04270 (LIC10895-96/LA3242) |  | TonB-dependent receptor | -3.262 | 2.32e-05 | 0.0355 |
| LIMLP_04275 (LIC10897/LA3241) |  | Hypothetical | -3.888 | 4.42e-05 | 0.00918 |
| LIMLP_04280 (LIC10898/LA3240) | *lipI48* | Hypothetical | -5.506 | 2.03e-13 | 0.00372 |
| LIMLP_08590 (LEPIC1767/LA2195)* |  | Hypothetical | -0.859 | 5.08e-08 |  |
| LIMLP_09650 (LIC11935/LA1968)* |  | Hypothetical | -1.787 | 3.26e-02 |  |
| LIMLP_14190 (LIC10709/LEPIN3051)** |  | Hypothetical lipoprotein | -0.679 | 1.88e-05 |  |
| LIMLP_14195 (LIC10708/LA3473)** |  | Hypothetical | -0.813 | 1.32e-12 |  |
| LIMLP_14200 (LIC10707/LA3474)** |  | Hypothetical GDSL-like lipase | -0.853 | 1.23e-05 |  |
| LIMLP_14205 (LIC10706/LA3475)** |  | Hypothetical lipoprotein | -0.847 | 4.75e-02 |  |
| LIMLP_14210 (LIC10705/LA3477) |  | Hypothetical lipoprotein | -0.915 | 3.05e-03 |  |
| LIMLP_14220 (LIC10703/LA3479) |  | Hypothetical | -0.793 | 1.13e-02 |  |
| LIMLP_14225 (LIC10702/LA3480) |  | Hypothetical | -0.775 | 2.14e-03 |  |
| LIMLP_15470 (LIC10454/LA3793) |  | Putative hemolysin | -2.154 | 3.32e-12 | 0.3041 |
| LIMLP_16720 (LIC13269/LA4102) | *vicK* | Response regulator | -1.611 | 5.80e-07 | 0.0752 |
| LIMLP_16725 (LIC13270/LA4104) | *vicR* | Signal transduction histidine kinase | -0.919 | 4.02e-03 | 0.4975 |
| LIMLP_18235 (LIC20078/LB099) |  | Hypothetical | -0.658 | 7.87e-03 |  |
| **Up-regulated genes** |  |  |  |  |  |
| LIMLP_02010 (LIC13086/LA3867)** |  | Hypothetical lipoprotein | 1.029 | 4.08e-02 |  |
| LIMLP_02795 (LIC12927/LA0666)* | *ccp* | Cytochrome C peroxidase | 2.773 | 8.69e-18 | 7.943 |
| LIMLP_05955 (LIC11219/LA2809)* | *ahpC* | Peroxiredoxin/alkylperoxidoreductase | 1.539 | 1.23e-05 | 2.01 |
| LIMLP_10145 (LIC12032/LA1859)* | *katE* | Catalase | 2.637 | 2.59e-24 | 4.897 |
| LIMLP_10150 (LIC12033/LA1858)* |  | Ankyrin repeat-containing protein | 2.867 | 4.65e-29 | 5.783 |

Significantly up-and down-regulated genes in the *perR* mutant with a Log<sub>2</sub>FC above 1 and below -0.5 and a p-value cutoff of 0.05.
<sup>a</sup> Gene numeration is according to Satou et al. (52). Corresponding genes of *L. interrogans* serovar lai strain 56601 and serovar Copenhageni Fiocruz strain L1-130 are indicated in parenthesis.
<sup>b</sup> Fold change in gene expression upon *perR* inactivation obtained by RT-qPCR experiments.
\* ORFs significantly up-regulated upon exposure to H<sub>2</sub>O<sub>2</sub> (Log<sub>2</sub>FC cutoff of 1).
\*\* ORFs significantly down-regulated upon exposure to H<sub>2</sub>O<sub>2</sub> (Log<sub>2</sub>FC cutoff of -1).

The *ank-katE* operon, encoded by LIMLP_10150-10145, *ahpC*, encoded by LIMLP_05955, and *ccp*, encoded by LIMLP_02795, were up-regulated upon *perR* inactivation. ChIP-PCR experiments showed that when *Leptospira* were cultivated in EMJH medium, PerR was bound to DNA fragments comprising the 25 to 191 bp upstream region to the *ank-katE* operon (Fig 4A). Lower non-significant PerR binding was detected inside the *ank-katE* operon (S5 Fig). This is consistent with a direct repression of the *ank-katE* operon by PerR. Significant PerR binding was observed from 150 to 350 bp upstream the LIMLP_02790, the ORF located immediately upstream the *ccp* ORF (Fig 4B and S5 Fig). ChIP experiments also showed a rather weak binding upstream the *ahpC* ORF (−298 to −123 region) (S5 Fig).

**Fig 4.**
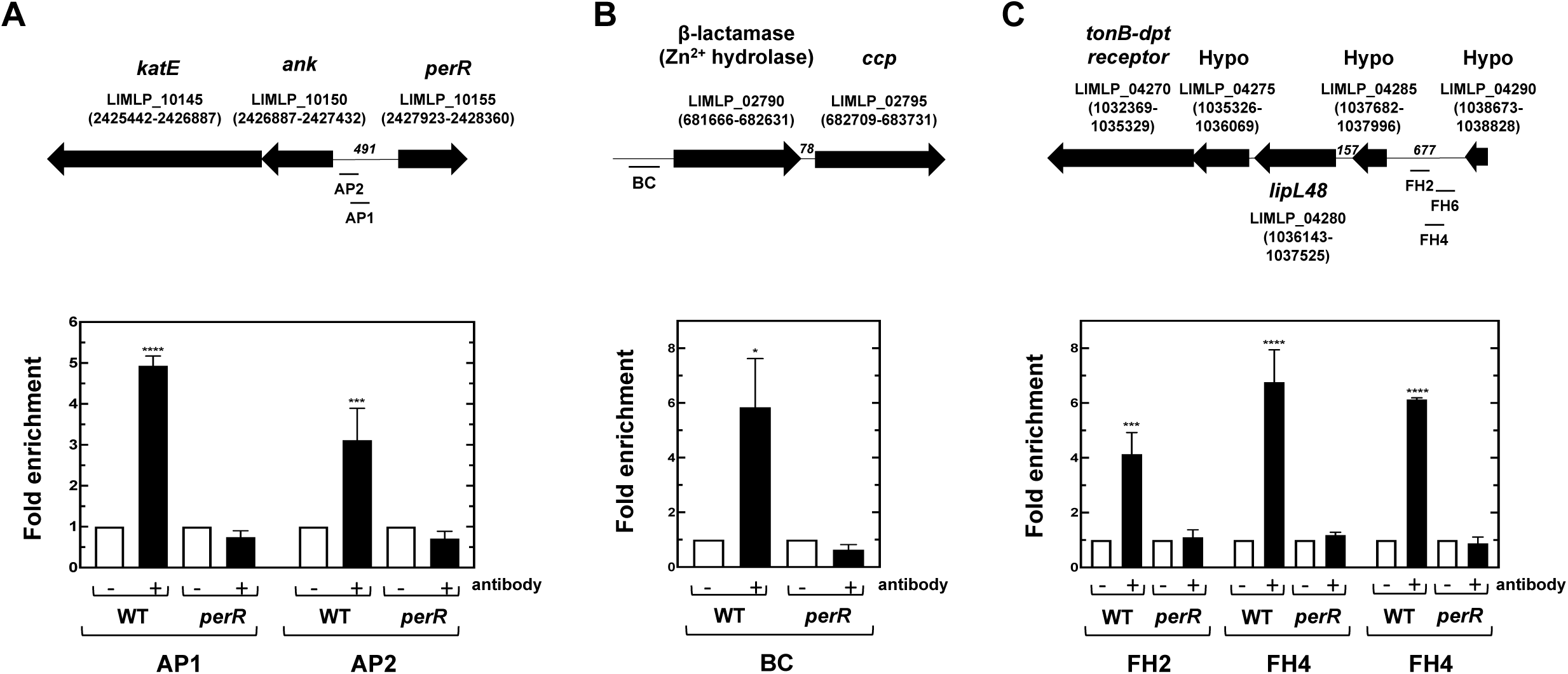
*In vivo* interaction between PerR and promoter regions of PerR-controlled genes. Chromatin immunoprecipitation was performed on *L. interrogans* WT and *perR* (M776) mutant strains in the presence or absence of the anti-PerR antibody as described in the Material and Methods section. Co-immunoprecipitated DNA fragments located in the *ank-katE* operon locus (A), in the *ccp* locus (B) and in the locus encoding a TonB-dependent transport system (C) were amplified by qPCR. The location of amplified fragments is indicated below the schematic representation of their respective locus. ORF location in the genome (according to Satou *et al*. (52)) is indicated into parenthesis and the length of intergenic regions are indicated in nucleotides in italic. Data are represented as fold enrichments and are means and SD of two independent biological replicates (****, adjusted p-value<0.0001; ***, adjusted p-value of 0.0006; *, adjusted p-value of 0.0422).

Determining PerR regulon has allowed the identification of genes whose expression is activated, directly or indirectly, by PerR (Table 3). A cluster composed of genes encoding a TonB-dependent transport (TBDT) system (LIMLP_04240-04255, encoding TonB, two ExbDs and ExbB, respectively) were dramatically down-regulated in the *perR* mutant (with a Log_2_FC of −5.47 to −4.60). The downstream TonB-dependent receptor- and LipL48-encoding ORFs were also down-regulated upon *perR* inactivation (with Log_2_FC of −3.26 and −5.50, respectively). LIMLP_04240 (*tonB*), LIMLP_04245 (*exbD*), and LIMLP_04250 (*exbD*) were organized as an operon and LIMLP_04270 (encoding a TonB-dependent receptor), LIMLP_04275, LIMLP_04280 (*lipl48*) and LIMLP_04285 constituted another operon (S6 Fig). ChIP-PCR assays indicated that a region upstream the LIMLP_04285-04270 operon (mapped from 436 to 617 bp) was significantly bound by PerR (Fig 4C) whereas lower binding was detected upstream the LIMLP_04265 ORF and within the LIMLP_04245 (S7 Fig).

A bicistronic operon composed of the response regulator VicR (LIMLP_16720) and the histidine kinase VicK (LIMLP_16725) of a two-component system (S8 Fig) was also down-regulated (with a Log_2_FC of −0.91 to −1.61, respectively) (Table 3).

Three ncRNAs were noticeably differentially expressed upon *perR* inactivation. The ncRNA rh859, located downstream *ccp* (LIMLP_02795), was up-regulated in the *perR* mutant (Log_2_FC of 2.50) as well as rh288 which overlaps one of the prophage locus ORF (LIMLP_00895). A 77 bp ncRNA (rh1263) located in the intergenic region upstream the TonB/ExbD_2_/ExbB-encoding operon (LIMLP_04255-04240) and downstream the ORF encoding a putative TonB-dependent receptor (LIMLP_04270) was significantly down-regulated in the *perR* mutant (Log_2_FC of −3.129).

Interestingly, among the PerR regulon, only genes whose expression is repressed by PerR were up-regulated when *Leptospira* were exposed to H_2_O_2_. Indeed, the expression of the *ank-katE* operon, *ahpC* and *ccp* were up-regulated in the *perR* mutant and in the presence of H_2_O_2_ whereas the expression of the ORFs encoding the TBDT, VicR and VicR was not dramatically or significantly altered by the presence of H_2_O_2_.

In order to determine the exact contribution of PerR in the gene expression increase upon exposure to H_2_O_2_ in *Leptospira*, the transcriptome of the *perR* mutant exposed to a sublethal dose of H_2_O_2_ was also obtained. The *ank-katE* operon, whose expression is directly repressed by PerR and increased in the presence of H_2_O_2_ in WT *Leptospira*, was not up-regulated in the presence of H_2_O_2_ when *perR* was inactivated (Fig 5). The amount of *ank-katE* operon expression in the *perR* mutant is in fact comparable to that in WT *Leptospira* exposed to a deadly dose of H_2_O_2_. This indicates that derepression of the *ank-katE* operon induced by the presence of H_2_O_2_ probably solely reflects PerR dissociation from DNA when PerR is oxidized. *AhpC* and *ccp* were still significantly up-regulated in the presence of H_2_O_2_ in the *perR* mutant (with Log_2_FC values of 2.298 and 1.874, respectively, see Fig 5). Therefore, an H_2_O_2_-induced mechanism increases the expression of these two genes even in the absence of PerR, even though their expression is repressed by this regulator.

**Fig 5.**
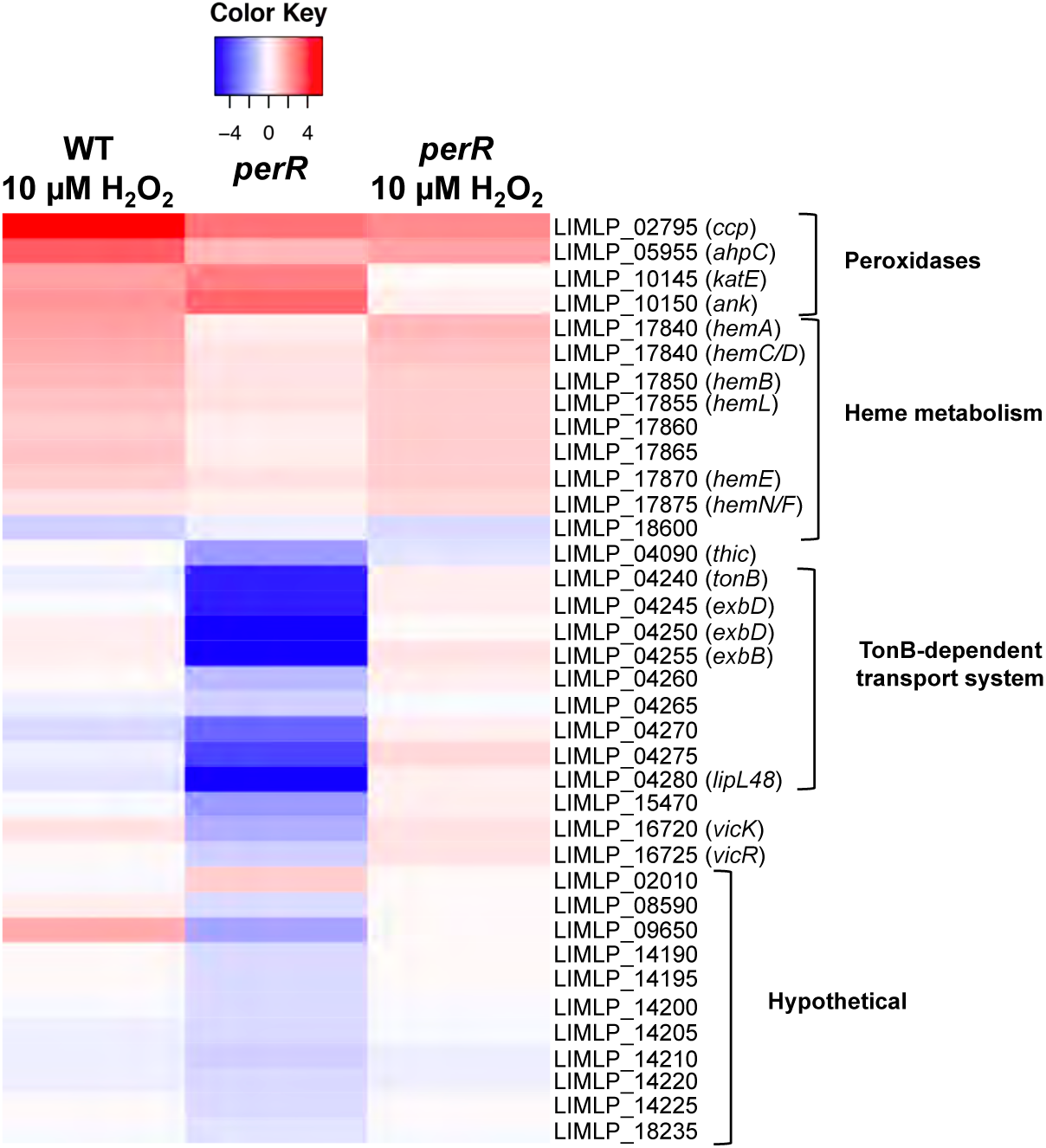
Comparison of differentially expressed genes upon exposure to hydrogen peroxide and *perR* inactivation. The expression of selected differentially-expressed genes determined by RNASeq when WT *L. interrogans* are exposed 30 min to 10 µM H_2_O_2_ was compared to that in the *perR* (M776) mutant exposed in the absence or presence of 10 µM H_2_O_2_. Genes are organized by their function, and their number and name are indicated on the right. The Heat Map color from blue to red indicates low to high Log_2_FC.

The expression of heme biosynthesis genes was not under the control of PerR and, as expected, their expression was still up-regulated in the *perR* mutant in the presence of H_2_O_2_ (Fig 5).

Interestingly, the ncRNA rh859, located downstream the *ccp*, was further up-regulated in the *perR* mutant upon exposure to H_2_O_2_ (Log_2_FC of 1.71). This indicates that the rh859 up-regulation induced by the exposure of WT *Leptospira* to H_2_O_2_ occurs to some extent independently of the presence of PerR. When the *perR* mutant was exposed to 10 µM of H_2_O_2_, 10 ncRNAs were significantly down-regulated with a log_2_FC below −2 whereas only one ncRNA was significantly down-regulated when WT cells were exposed to 10 µM of H_2_O_2_. For several of them (rh96, rh367, rh753, rh928, rh2114, rh2850, rh4234, and rh4918), such a down-regulation was not observed upon exposure of WT cells to 10 µM H_2_O_2_ and only observed in the absence of PerR.

Altogether, these findings indicate that not all H_2_O_2_-regulated genes belong to the PerR regulon in pathogenic *Leptospira* and several PerR-regulated genes were not regulated by H_2_O_2_ (Fig 5). Likewise, differential expression of putative ncRNAs upon exposure to hydrogen peroxide is not necessarily exerted by PerR.

### Role of the PerR-regulated genes in defenses against ROS and virulence in *Leptospira*

RNASeq experiments have allowed the identification of differentially expressed ORFs in the presence of peroxide and/or upon *perR* inactivation. These ORFs might encode cellular factors required for the adaptation of pathogenic *Leptospira* to ROS and an important question is to experimentally establish and understand the role of this factors in this adaptation.

Genetic manipulation of pathogenic *Leptospira* is still a challenge and functional studies in these bacteria mainly relies on random insertion transposon. Our laboratory has constructed a transposon mutant library (30) and several mutants inactivated in differentially-expressed ORFs upon exposure to H_2_O_2_ or upon *perR* inactivation were available in our library.

Catalase, AhpC, and CCP were the peroxidases up-regulated in the presence of H_2_O_2_ and repressed by PerR. Only *katE* and *ahpC* mutants were available in the transposon mutant library and we have studied the ability of these mutants to grow in the presence of H_2_O_2_ and paraquat, a superoxide-generating compound. These two mutants had a comparable growth rate in EMJH medium (Fig 6A) but when the medium was complemented with 0.5 mM H_2_O_2_, the ability of the *katE* mutant to divide was dramatically impaired (Fig 6B). The growth rate of the *ahpC* mutant in the presence of H_2_O_2_ was comparable to that of the WT strain (Fig 6B). When the EMJH medium was complemented with 2 *μ*M paraquat, the growth of the *ahpC* mutant was considerably reduced, indicating a high sensitivity to superoxide (Fig 6C).

**Fig 6.**
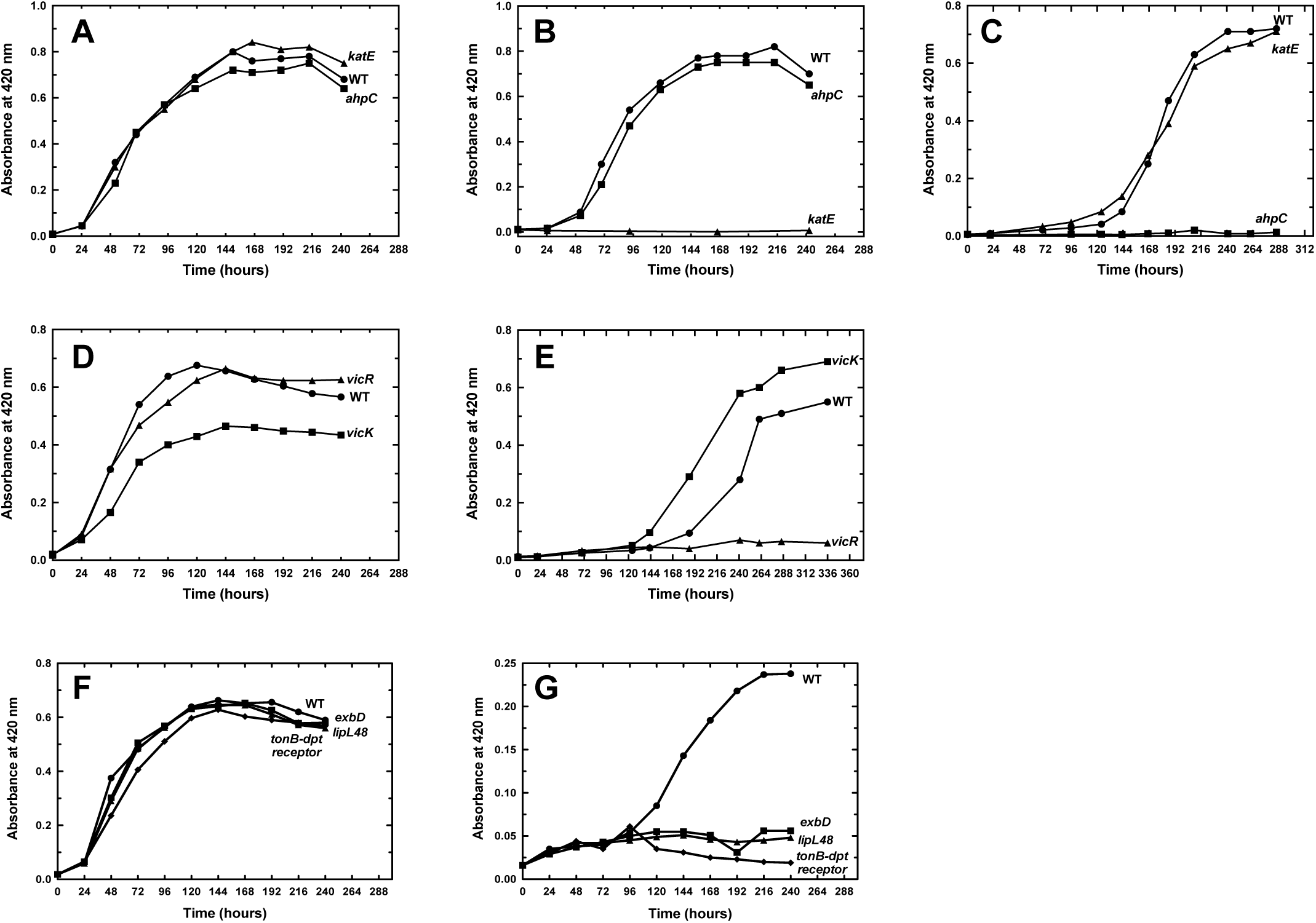
Effect of the inactivation of PerR-controlled genes on *Leptospira* growth in the presence of ROS. *L. interrogans* WT, *katE* (Man69), *ahpC* (Man1368), *vicK* (Man1448), *vicR* (Man899), *tonB-dpt receptor* (Man1022), *exbD* (Man782), and *lipl48* (Man1089) mutant strains were cultivated in EMJH medium (A, D, F) or in the presence of 2 mM H_2_O_2_ (B) or of 2 µM paraquat (C, E, G). Growth was assessed by measure of absorbance at 420 nm. Data are representative of at least four independent replicates.

In other bacteria including *E. coli* and *B. subtilis, katE* is produced in higher amount during stationary phase (31,32), and in order to further characterized the role of *katE* in *Leptospira* survival under oxidative stress, we investigated the survival of stationary phase-adapted *Leptospira* in the presence of H_2_O_2_. *L. interrogans* WT cells were cultivated in EMJH medium and samples were harvested in the logarithmic phase (Fig 7A, sample 1), at the entry in stationary phase (Fig 7A, sample 2) and in advanced stationary phase (Fig 7A, sample 3).

**Fig 7.**
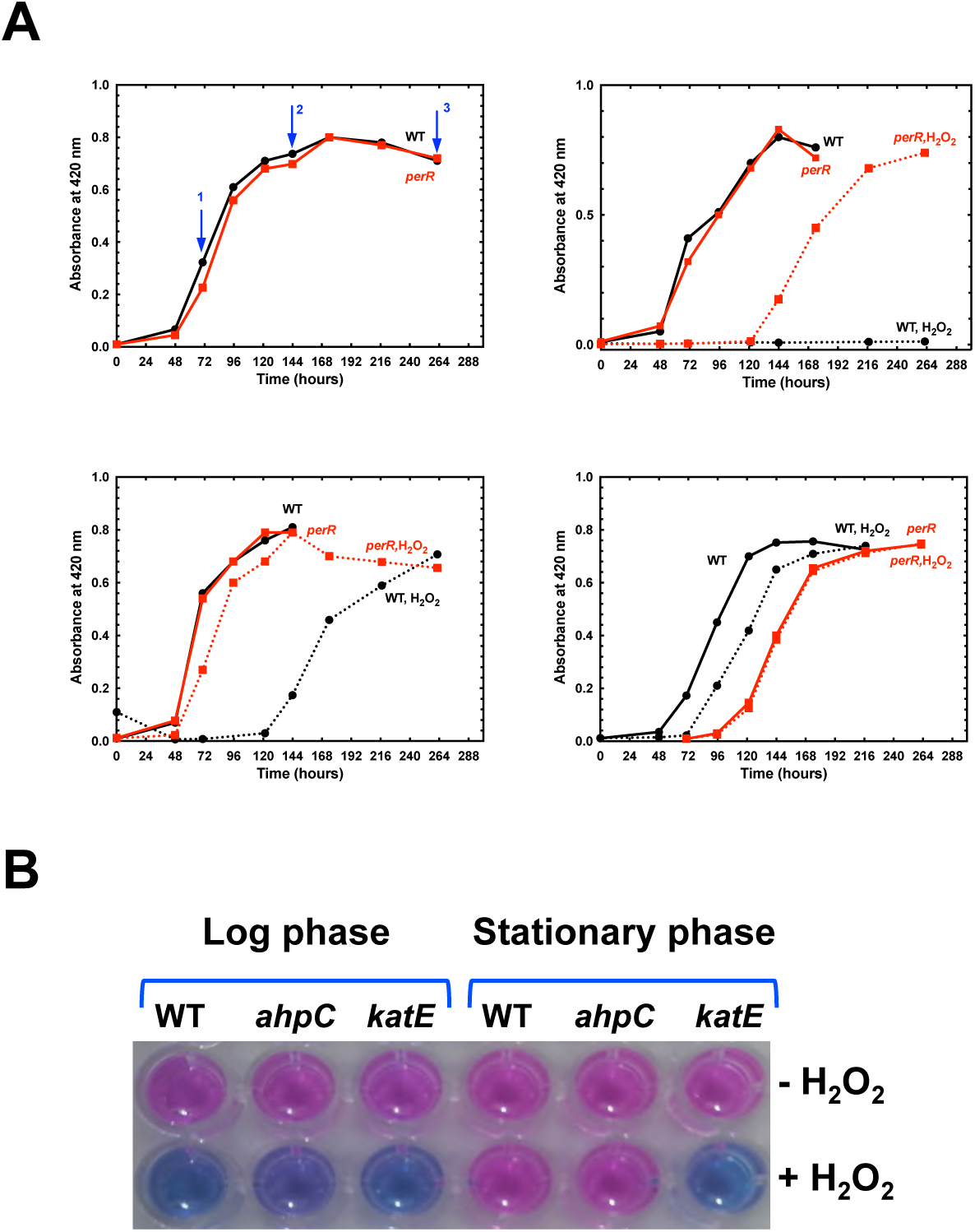
Role of catalase and AhpC in the stationary phase-adapted *Leptospira* tolerance to hydrogen peroxide. (A) *L. interrogans* WT (black line) and *perR* mutant (M776) (red line) strains were cultivated in EMJH medium and samples were taken at the exponential phase (at OD_420_ nm ≈ 0.3, left upper panel, blue arrow 1), at the entry of stationary phase (atOD_420_ nm ≈ 0.7, left upper panel, blue arrow 2), and at advanced stationary phase (at OD_420_ nm ≈ 0.7, 5 days after the entry in stationary phase, left upper panel, blue arrow 3) and used to inoculate a new EMJH medium in the absence (plain line) or presence of 2 mM H_2_O_2_ (dashed line). The growth curve with samples taken in the exponential phase (samples 1), in the entry of stationary phase (samples 2) and at advanced stationary phase (samples 3) are represented in the right upper, the left lower, and the right lower panels, respectively. (B) *L. interrogans* WT, *katE* (Man69) and *ahpC* (Man1368) mutant strains were cultivated in EMJH medium until the exponential or stationary phases and incubated for 30 min in the absence or presence of 10 mM H_2_O_2_. Cell viability was assessed by the ability of the cells to reduce the blue rezasurin into a pink resorufin using the Alamar Blue assay as described in the Material and Methods section.

Each sample was used to inoculate a new batch of EMJH medium in the absence or presence of 2 mM H_2_O_2_. As seen in Fig 7A, when EMJH was inoculated with *Leptospira* cells at logarithmic phase, *Leptospira* were not able to divide in the presence of 2 mM H_2_O_2_. However, when the culture medium was inoculated with *Leptospira* cells at the beginning of the stationary phase, *Leptospira* acquired a greater resistance to 2 mM H_2_O_2_ as seen by their ability to grow (Fig 7A). An even higher ability to grow in the presence of a deadly dose of H_2_O_2_ was observed when the EMJH medium was inoculated with *Leptospira* at advanced stationary phase (Fig 7A). This indicates that *Leptospira* cells acquire a higher tolerance to hydrogen peroxide at stationary phase. Interestingly, this acquired tolerance to H_2_O_2_ was independent of PerR since the *perR* mutant also acquired a higher ability to grow in the presence of 2 mM H_2_O_2_ when at stationary phase (Fig 7A). In order to determine which peroxidase was responsible for this acquired tolerance to H_2_O_2_, the survival of WT, *ahpC* and *katE* mutant strains was tested in logarithmic phase and was compared with that in stationary phase. As seen in Fig 7B, a 30 min exposure to 10 mM H_2_O_2_ led to dramatic loss of survival of all strains at logarithmic phase. WT and *ahpC* mutant strains were able to acquire a higher resistance to H_2_O_2_ when placed at stationary phase whereas the *katE* mutant did not. Therefore, *katE* is essential for the stationary phase-acquired resistance to H_2_O_2_ and this probably involves another regulation mechanism than that exerted by PerR.

Among the genes repressed by PerR, only mutants inactivated in LIMLP_04245 (*exbD*), LIMLP_04270 (*tonB-dpt receptor*), LIMLP_04280 (*lipl48*), LIMLP_16720 (*vicR*), and LIMLP_16725 (*vicK*), were available in the transposon mutant library. All these mutants but *vicK* had a growth comparable to that of the WT strain in EMJH medium (Figs 6D and 6F). Despite the fact that *vicK* had a reduced ability to divide in EMJH medium, this mutant strain had a slightly greater resistance to 2 *μ*M paraquat than that of the WT (Fig 6E). In the same condition, the *vicR, exbD, tonB-dpt receptor* and *lipl48* mutant strains had a lower ability to grow than the WT strain (Figs 6E and 6G). Altogether, these findings suggest that some of the PerR-repressed ORF are involved in *Leptospira* defense against superoxide.

Catalase has been shown to be essential for *Leptospira* virulence (9). We investigated whether other PerR-controlled genes were also required for *Leptospira* virulence. The different mutants were used in infection experiments in the acute model for leptospirosis. *VicK, exbD*, and *lipl48* mutants did not exhibit dramatic altered virulence when 10^6^ bacteria were injected peritoneally in hamsters (Figs 8A and 8B). In order to further challenge the role of the TonB-dependent transport system in *Leptospira* virulence, we tested whether a lower dose of infection with the *tonB-dpt receptor* and *exbD* mutants would result in a virulence attenuation. As seen in Fig 8C, when 10^4^ bacteria were injected peritoneally in hamsters, animals infected with the *exbD* mutant exhibited 25% survival at 32 days post infection with no sign of leptospirosis. However, this slight virulence attenuation was not considered as statistically significant. Therefore, inactivation of the TonB-dependent transport system or of the two-component system VicKR does not have a drastic consequence on *Leptospira* virulence in our acute model of infection.

**Fig 8.**
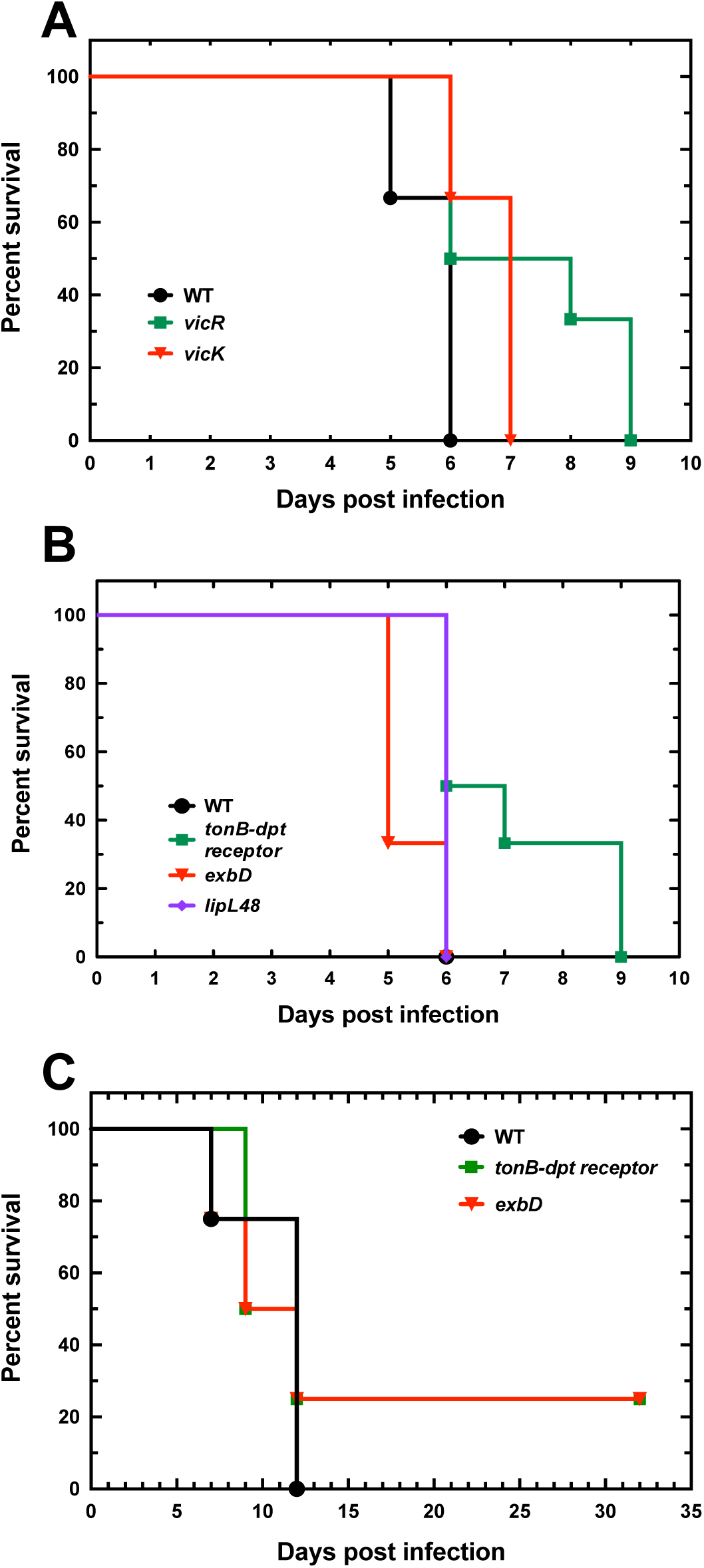
Role of PerR-controlled ORFs in *Leptospira* virulence. 10^6^ of WT, *vicK* (Man1448) and *vicR* (Man899) mutant strains (A), or the *tonB-dpt receptor* (Man1022), *exbD* (Man782), *lipl48* (Man1089) mutant strains (B) or 10^4^ of WT, *exbD* (Man782), or the *tonB-dpt receptor* (Man1022) mutant strains (C) were injected intraperitoneally in hamsters (n=4-8) as described in Material and Methods section.

## Discussion

Reactive oxidative species are powerful and efficient weapons used by the host innate immunity response to eliminate pathogens. The ability of pathogenic *Leptospira* to detoxify hydrogen peroxide, one of the ROS produced upon *Leptospira* infection and pathogenicity, is essential for these pathogenic bacteria virulence (9). Because *Leptospira* are also environmental aerobic bacteria, they will also face low concentrations of ROS endogenously produced through the respiratory chain or present in the outside environment. The present study has used RNASeq technology to determine the adaptive response of pathogenic *Leptospira* to hydrogen peroxide. Our study allowed, for the first time, a global identification of all the cellular factors solicited by pathogenic *Leptospira* to adapt to the presence of H_2_O_2_. *L. interrogans* were subjected to two different treatments. A short exposure in the presence of a sublethal dose of hydrogen peroxide (30 min. with 10 *μ*M H_2_O_2_) and a longer exposure with a lethal concentration of hydrogen peroxide (60 min. with 1 mM H_2_O_2_), that could mimic the hydrogen peroxide concentrations encountered inside a host. Our findings indicate that the peroxidic stress response is timely-orchestrated and dose-dependent. *L. interrogans* can sense and rapidly respond to H_2_O_2_ concentrations as low as 10 *μ*M by up-regulating the catalase (encoded by *katE*) and two peroxidases, an AhpC and a CCP (Fig. 9A). Heme biosynthesis-encoding genes were also up-regulated probably because catalase and CCP have heme-dependent peroxidase activities. These three peroxidases are the first-line of defense allowing detoxification of H_2_O_2_, and among these three enzymes, *katE*-encoded catalase has a major role in protecting *L. interrogans* from the deadly effect of hydrogen peroxide, during logarithmic phase but also during stationary phase. Arias et al. (21) showed that *E. coli* cells overexpressing the *L. interrogans* AhpC displayed a higher survival in the presence of H_2_O_2_ and *tert*-butylhydroperoxide and recombinant AhpC was able to reduce these peroxides using the *E. coli* thioredoxin system as a cofactor. In our study, an *ahpC* mutant did not exhibit an altered tolerance toward H_2_O_2_; instead, this mutant had a lower ability to grow in the presence of paraquat, a superoxide-generating chemical. Although we cannot rule out that the inactivation of *ahpC* triggers an increase in catalase activity to compensate the absence of AhpC, our findings might indicate a role of this peroxidase in detoxification of superoxide or of H_2_O_2_ produced from the catabolism of superoxide. The role of CCP in degrading H_2_O_2_ in pathogenic *Leptospira* has never been investigated. Whether CCP fulfills such a role or whether CCP rather acts as an electron acceptor for the respiratory chain, as demonstrated in *E. coli* (33), will require obtaining a deletion mutant by allelic exchange since a *ccp* mutant was not available in the transposon mutant library.

**Fig 9.**
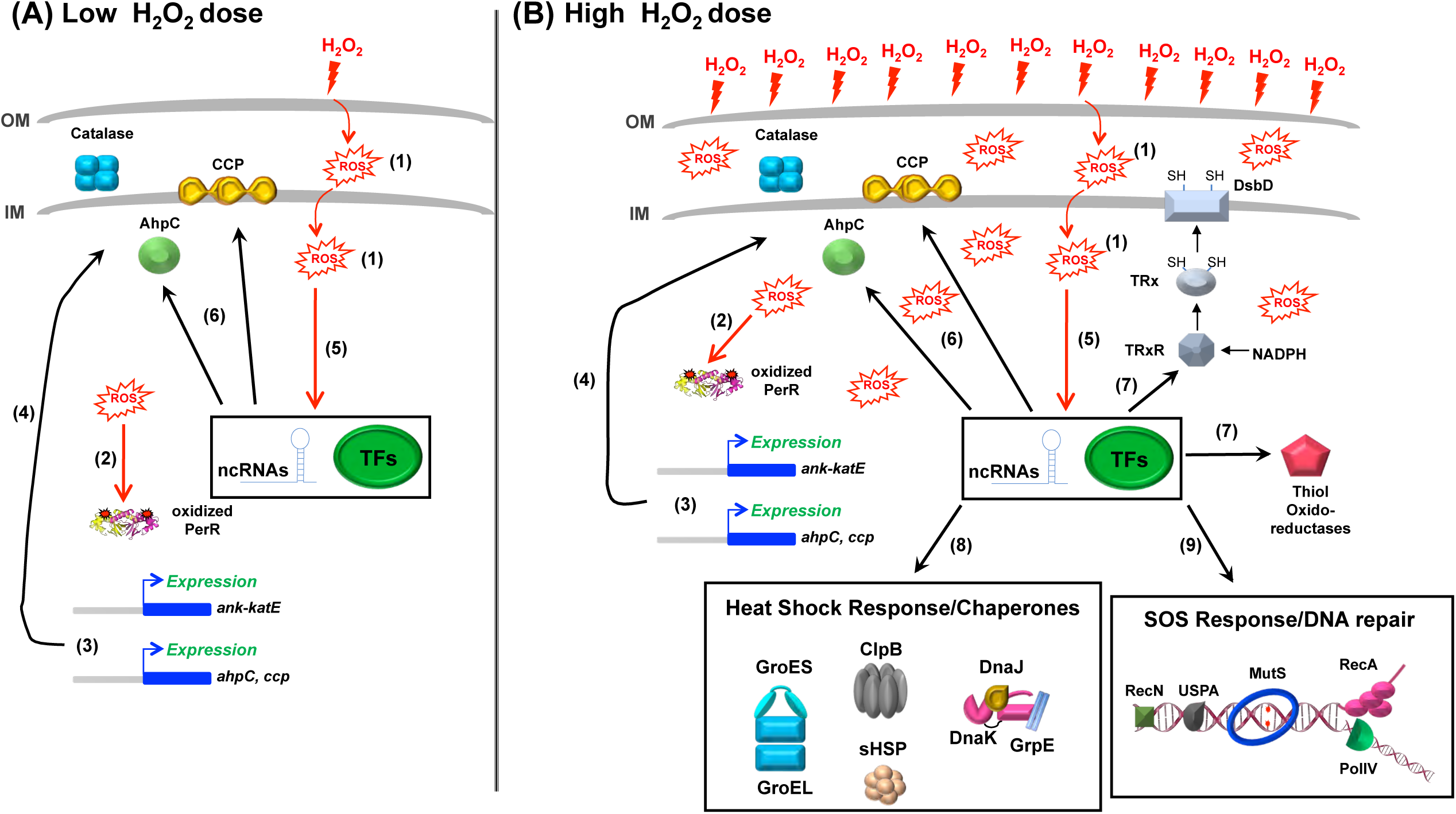
Schematic representation of the pathogenic *Leptospira* adaptive response to hydrogen peroxide. In the presence of low H_2_O_2_ dose (A), ROS are produced in the cell (1), PerR is oxidized (2) and dissociates from DNA regions in the locus of the three peroxidase-encoding genes (*ank-katE* operon, *ahpC, ccp*), leading to their derepression (3) and increased production of catalase, AhpC and CCP (4). Other transcriptional regulator (TFs) and non-coding RNAs whose expression is affected in the presence of ROS (5) probably participate in the H_2_O_2_-induced increase of AhpC and CCP production (6). As a result, the activities of catalase, AhpC and CCP allow maintaining ROS at a harmless level. The increased expression of heme biosynthesis genes, which is PerR-independent and probably participated in the peroxidase activities of catalase, AhpC and CCP, is not represented here. When the level of ROS overwhelms the detoxification capacity of the up-regulated peroxidases and becomes damaging for the cellular constituents (B), in addition of a higher production of catalase, AhpC and CCP, other machineries are up-regulated such as thiol oxido-reductases (including thioredoxin, DsbD, etc.) (7), molecular chaperones (8) and DNA repair proteins (9). The increased expression of the aforementioned machineries is PerR-independent and probably involves other transcriptional regulator (TFs) and noncoding RNAs.

The up-regulation of catalase, AhpC and CCP is probably sufficient to rapidly degrade H_2_O_2_ and avoid accumulation of ROS inside the cells. However, when H_2_O_2_ reachs a level that overwhelms the H_2_O_2_ detoxification machinery, as occurred when *L. interrogans* are exposed to 1 mM H_2_O_2_ (Fig 9B), not only *L. interrogans* solicited the aforementioned peroxidases but additional enzymes with a putative role as antioxidants and/or in repair of oxidized cysteines in proteins were also up-regulated, including several thiol oxidoreductases (thioredoxin, glutaredoxin, DsbD and Bcp-like proteins). The induction of several genes of the LexA regulon (*recA, recN, dinP*) and other genes with putative role in DNA repair (*mutS, radC*) suggests that these concentration of H_2_O_2_ induced oxidative damage to DNA and a need for the SOS response. Surprisingly, the classical repair mechanism for oxidized methionine residues (such as methionine sulfoxide reductases) or damages to iron-sulfur clusters in proteins (the Suf machinery) were not more expressed in the presence of H_2_O_2_ as if this repair mechanisms were not required under such oxidative damage-inducing condition. Also, the redox-regulated chaperone Hsp33 involved in protecting form aggregation and promoting the refolding of oxidatively-damaged proteins, was not up-regulated. Instead, canonical ATP-dependent (DnaK/J/GrpE, GroEL/ES, ClpB) and -independent (small Hsps) molecular chaperones were dramatically more expressed, suggesting that 1 mM H_2_O_2_ results in protein aggregation and unfolding.

The up-regulation of all these detoxification and repair mechanisms correlated with a down-regulation of genes encoding transcription and translation factors, protein secretion, *Leptospira* motility and chemotaxis, as well metabolism pathways including ATP and cobalamin (vitamin B12) biosynthesis, that might explain the slowdown in growth induced by the presence of H_2_O_2_.

Several of the factors whose expression is up-regulated upon exposure to H_2_O_2_ were also up-regulated when *Leptospira* are cultivated in DMCs implemented in rat peritoneal cavities (16). Among those were the peroxidases AhpC and CCP and their repressor PerR. Also, *rcc1*, one of the highest up-regulated gene upon exposure to 1 mM H_2_O_2_, as well as genes encoding DNA repair proteins (LIMLP_11400, LIMLP_16520, and LIMLP_16525), were also up-regulated in the DMC model. This might indicate that the oxidative stress condition used in the present study properly reproduces that encountered by *Leptospira* during infecting a mammalian host. Many ORFs of the H_2_O_2_ adaptive response identified in this study have been shown to be also up-regulated upon other host-related conditions such as at the host temperature of 37°C (Catalase, GroEL/ES, DnaK/J/GrpE, small HSPs, ClpB, RadC) (34–36), host osmolarity (RadC, the LIMLP_16520-encoded DNA repair exonuclease, DsbD and the LIMLP_00770-encoded dithiol disulfide isomerase) (37), or under iron-limited condition as encountered inside a host (such as ORFs encoding TonB-dependent receptors (LIMLP_14160 and LIMLP_08410), Imelysin (LIMLP_14180), the lipoprotein LruB (LIMLP_14170)) (10). Therefore, the H_2_O_2_ adaptive response overlaps to some extent with other stress responses. In fact, the accumulation of oxidatively-damaged proteins and DNAs could trigger a general stress response. The change in expression of other stress-related regulators such HrcA and LexA suggests that the presence of ROS elicits heat shock and SOS responses.

Comparing the H_2_O_2_-induced changes in gene expression in the *perR* mutant with that in WT cells, indicated that PerR contributes only partially to the H_2_O_2_-induced gene regulation. Among the genes whose expression is markedly changed upon exposure to H_2_O_2_, only *katE, ahpC* and *ccp* are under the control of PerR. Surprisingly, even in the absence of PerR, *ahpC* and *ccp* expressions are still increased upon exposure to H_2_O_2_, suggesting that additional regulatory mechanisms are involved in the H_2_O_2_-induced gene regulation. In fact, several genes encoding transcriptional regulators, two component systems, and sigma factors had their expression altered by the presence of H_2_O_2_, corroborating the involvement of other regulators in the adaptive response to oxidative stress in pathogenic *Leptospira* (Fig. 9B). Moreover, we have identified several ncRNAs that might also influence the expression of the H_2_O_2_-regulated genes. For instance, many ncRNAs with increased or reduced expression upon *Leptospira* exposure to H_2_O_2_ are located in the vicinity of ORFs with increased or reduced expression in the same condition. Noticeably, rh859 located downstream *ccp* might participate in the increased expression of this gene, together with the derepression induced by PerR dissociation from DNA in the presence of H_2_O_2_ (Fig. 9). Also, the ncRNA rh3130 might be involved in the up-regulation of the sHsp-encoding operon upon exposure to H_2_O_2_. Therefore, our study has unveiled the complexity of the regulatory network involved in the leptospiral adaptive response to oxidative stress.

Comparison of the transcriptome of the *perR* mutant determined in this study with that determined previously by Lo *et al.* (10) pinpoints several discrepancies. For instance, our study did not demonstrate that heme biosynthesis genes are under the control of PerR and the expression of *ahpC* was not affected in the *perR* mutant in the study of Lo *et al.* (10). These contradictions can be explained by the experimental conditions used to determine the transcriptome of the *perR* mutant in this previous study which, in fact, has compared WT cells cultivated in EMJH medium with *perR* mutant cells cultivated in EMJH medium in the presence of kanamycin. Due to the relation between antibiotic and oxidative stresses, the presence of an antibiotic might have influenced the expression of ROS-related genes, such as heme genes or *ahpC*.

In the present study, we have further studied the PerR-mediated gene expression control by investigating the direct binding of PerR to target genes. We have shown that PerR binds the upstream region of the *ank-katE* operon, indicating that PerR directly represses this operon. In our study, *Leptospira* PerR was shown to bind more than 1 kb upstream *ccp*. Such a binding at a distal site from the *ccp* promoter region would be consistent with a control of *ccp* expression mediated by DNA deformation (such as looping) induced by PerR binding. Also, our data suggest the involvement of a ncRNA located immediately downstream *ccp* participating, together with PerR, in the control of *ccp* expression. Likewise, PerR was shown to bind in the intergenic region of LIMLP_04285 and LIMLP_04290, at a distal site (500 bp) upstream of the LIMLP_04285 ORF. In addition, two putative ncRNAs were identify in this locus (rh1269 and rh1263). Whether PerR binding at a distal site from the promoter regions and the presence of these ncRNAs participate in the control of the expression of this gene cluster will need to be demonstrated. The use of the ChIP-Seq approach will be pivotal for achieving this aim.

Among the ORF that are significantly up-regulated in the presence of H_2_O_2_, catalase and ClpB have been shown to be required for *Leptospira* survival under oxidative stress and virulence (9,36). In the present study, we have confirmed the essential role of *katE* for the defense against H_2_O_2_, particularly in stationary phase. Furthermore, we have identified new ORFs that participate in *Leptospira* survival in the presence of ROS. Indeed, our findings indicate that the AhpC peroxidase, a TonB-dependent transport system (TBDT), the lipoprotein LipL48, and the response regulator VicR are involved in *Leptospira* survival in the presence of a superoxide-generating compound. Interestingly, pathogenic *Leptospira* do not encode any genes homolog to a superoxide dismutase or reductase, nor they exhibit any SOD activity (38). This is quite intriguing as it is generally believed that all aerobic bacteria do have a SOD. One fundamental question is to understand the mechanism these pathogenic bacteria use to detoxify superoxide produced endogenously during the respiratory chain or exogenously by phagocytic cells during infection. Our study is the first to identify cellular factor in pathogenic *Leptospira* involved in survival in the presence of superoxide-generating compound. AhpC could detoxify H_2_O_2_ produced upon the reduction of superoxide, but the exact function of ExbD, the TBDT, and LipL48 in superoxide detoxification is still unclear. In bacteria, ExbD is part of inner membrane complex TonB/ExbD/ExbB that uses proton motive force to provide the energy necessary by TBDT for uptake of an iron source. The presence of LipL48-encoding ORF in the same operon as the TBDT strongly suggests that these two proteins are functionally linked. This TBDT machinery could be solicited for iron uptake when iron concentration inside *Leptospira* is low as a result of the presence of ROS. Indeed, one can imagine that *Leptospira* lower the level of free intracellular iron to prevent worsening the production of ROS inside the cells by the Fenton reaction. Consistent with this is the down-regulation of some of the gene encoding this TBDT machinery upon exposure to H_2_O_2_ (Fig. 1). Alternatively, the TBDT machinery could also be involved in the uptake of another metal (such as zinc or manganese) that could be used by a ROS detoxification enzyme or even act by itself as ROS scavenger. Indeed, manganese has been shown to scavenge superoxide in *Lactobacillus plantarum* and *Neisseria gonorrhoeae*, independently to any SOD activity (39,40).

None of the mutants inactivated in these ORF exhibited a dramatic reduction in virulence, suggesting that these mechanisms do not have a pivotal role in *Leptospira* during infection or that redundant activities compensate for their absence. Experiments using other infection routes (ocular or subcutaneous routes) might result in different outcomes.

In conclusion, the present study has revealed, for the first time, the genome-wide general adaptive response to peroxide in pathogenic *Leptospira*, unfolding putative biological pathways *Leptospira* have evolved to overcome the deadly effect of ROS. Peroxide adaptive response involves detoxifying enzymes, molecular chaperones, DNA repair machinery, and transporters. This adaptive response also engages a large number of non-annotated and sometimes *Leptospira* specific ORFs reflecting the submerged part of the iceberg in these bacteria physiology. We have also uncovered a regulatory network of transcriptional regulators, sigma factors, two component systems and non-coding RNA that orchestrate together with PerR the peroxide adaptive response.

## Materials and Methods

### Bacterial strains and growth condition

*L. interrogans* serovar Manilae strain L495 and transposon mutant strains (see S3 Table for a complete description of the transposon mutants used in this study) were grown aerobically at 30°C in Ellinghausen-McCullough-Johnson-Harris medium (EMJH) (41) with shaking at 100 rpm. Cell growth was followed by measuring the absorbance at 420 nm.

### RNA purification

Virulent *L. interrogans* serovar Manilae strain L495 and *perR* mutant M776 with less than three *in vitro* passages were used in this study. Four independent biological replicates of exponentially grown WT and *perR* mutant *L. interrogans* strains were incubated in the presence or absence of 10 µM H_2_O_2_ for 30 min at 30°C. WT L495 strain was also incubated in the presence of 1 mM H_2_O_2_ for 60 min at 30°C. Harvested cells were resuspended in 1 ml TRIzol™ (ThermoFisher Scientific) and stored at −80°C. Nucleic Acids were extracted with chloroform and precipitated with isopropanol as described earlier (15). Contaminating genomic DNA was removed by DNAse treatment using the RNAse-free Turbo DNA-free™ turbo kit (ThermoFisher Scientific) as described by the manufacturer. The integrity of RNAs (RIN > 7.6) was verified by the Agilent Bioanalyzer RNA NanoChips (Agilent technologies, Wilmington, DE).

### RNA Sequencing

rRNA were depleted from 0.5 µg of total RNA using the Ribo-Zero rRNA Removal Kit (Bacteria) from Illumina. Sequencing libraries were constructed using the TruSeq Stranded mRNA Sample preparation kit (20020595) following the manufacturer’s instructions (Illumina). The directional libraries were controlled on Bioanalyzer DNA1000 Chips (Agilent Technologies) and concentrations measured with the Qubit® dsDNA HS Assay Kit (ThermoFisher). Sequences of 65 bases were generated on the Illumina Hiseq 2500 sequencer.

Bioinformatics analyses were performed using the RNA-seq pipeline from Sequana (42). Reads were cleaned of adapter sequences and low-quality sequences using cutadapt version 1.11 (43). Only sequences of at least 25 nt in length were considered for further analysis. Bowtie version 1.2.2 (44), with default parameters, was used for alignment on the reference genome (*L. interrogans* serovar Manilae strain UP-MMC-NIID LP, from MicroScope Platform). Genes were counted using featureCounts version 1.4.6-p3 (45) from Subreads package (parameters: -t gene -g locus_tag -s 1).

Count data were analyzed using R version 3.5.1 (46) and the Bioconductor package DESeq2 version 1.20.0 (47). The normalization and dispersion estimation were performed with DESeq2 using the default parameters and statistical tests for differential expression were performed applying the independent filtering algorithm. A generalized linear model including the replicate effect as blocking factor was set in order to test for the differential expression between *Leptospira* samples. Raw p-values were adjusted for multiple testing according to the Benjamini and Hochberg (BH) procedure (48) and genes with an adjusted p-value lower than 0.005 and a Log_2_FC higher than 1 or lower than −1 were considered differentially expressed. The Fisher statistical test was used for the COG (Clusters of Orthologous Groups) classification.

### Quantitative RT-qPCR experiments

cDNA synthesis was performed with the cDNA synthesis kit (Biorad) according to the manufacturer’s recommendation. Quantitative PCR was conducted with the SsoFast EvaGreen Supermix (Biorad) as previously described (9,15). Gene expression was measured using *flaB* (LIMLP_09410) as a reference gene.

### Non-coding RNA identification

Sequencing data from the *Leptospira* WT and *perR* mutant strains incubated in the absence or presence of H_2_O_2_ were processed with Trimmomatic (49) to remove low-quality bases and adapter contaminations. BWA mem (version 0.7.12) was used to discard the reads matching *Leptospira* rRNA, tRNA or polyA sequences and to assign the resulting reads to *Leptospira* replicons. Then Rockhopper (50) was used to re-align reads corresponding to separate replicons and to assemble transcripts models. The output was filtered to retain all transcripts longer than 50 nucleotides not overlapping within 10 nucleotides with NCBI annotated genes on the same orientation, and showing a minimum Rockhopper raw count value of 50 in at least two isolates. This high-quality set of 778 new sRNA was subjected to differential expression analysis with Rockhopper, adopting a Benjamini-Hochberg adjusted P-value threshold of 0.01. For each non-coding RNAs, putative function was identified by BLAST using the Rfam database (51).

### ChIP-qPCR

Chromatin immunoprecipitation was performed by incubating exponentially growing *Leptospira* WT or *perR1* mutant cells 40 min with 1% formaldehyde at 30°C. The reaction was stopped by the addition of 400 mM glycine. Cells were then washed with TBS buffer and resuspended in buffer A (50 mM HEPES-KOH pH7.5, 150 mM NaCl, 1 mM EDTA, 1% Triton X-100) containing a protease inhibitor cocktail. Cells were sonicated 7 cycles of 15 min. and centrifuged. The supernatant was incubated 3 hours at 4°C with 50 µl of washed Dynabead Pan rabbit IgG. The samples were incubated in the absence or in the presence of anti-PerR serum (at a dilution of 1:750) for 2 hours at 4°C. The samples were successively washed with buffer A containing 500 mM NaCl, with buffer B (10 mM Tris-HCl pH8, 1 mM EDTA, 0.1% Nonidet-P40, 0.5% sodium deoxycholate) and with buffer C (10 mM Tris-HCl pH7.5, 1 mM EDTA). The elution was performed with 100 µl of elution buffer (50 mM Tris-HCl pH7.5, 10 mM EDTA, 1% SDS, 150 mM NaCl, 0.5% Triton X-100) upon an ON incubation at 37°C. An incubation with protease K (2 hours at 65°C) allowed elimination of proteins and DNA fragments were purified. DNA fragments were amplified by qPCR using the SsoFast EvaGreen Supermix (Biorad). Results were normalized by the Fold enrichment method (signal over background) calculated using the following formula: 2^ΔΔCq (where ΔΔCq is Cq_with antibody_ – Cq_without antibody_).

### Determination of cell viability

Cell survival was determined by incubating exponentially growing *L. interrogans* cells (≈ 10^8^/ml) in EMJH in the presence or absence of H_2_O_2_. Cells were then incubated with rezasurin (Alamar Blue® Assay, ThermoFisher Scientific) for 24h. Viability is assessed by the reduction of blue resazurin into pink resorufin. Plating experiments were performed by diluting treated and non-treated cells in EMJH in the absence of H_2_O_2_ and plating the samples on EMJH agar medium. Colonies were counted after one month incubation at 30°C.

### Infection experiments

*L. interrogans* WT and mutant strains were cultivated in EMJH medium until the exponential phase and counted under a dark-field microscope using a Petroff-Hauser cell. 10^4^ or 10^6^ bacteria (in 0.5 ml) were injected intraperitoneally in groups of 4-8 male 4 weeks-old Syrian Golden hamsters (RjHan:AURA, Janvier Labs). Animals were monitored daily and sacrificed when endpoint criteria were met (sign of distress, morbidity). The protocol for animal experimentation is conformed to the guidelines of the Animal Care and Use Committees of the Institut Pasteur (CETEA #2016-0019).

## Acknowledgement

We would like to thank Maya Long and Clémence Mouville for their excellent and efficient technical help. Crispin Zavala-Alvarado is part of the Pasteur – Paris University (PPU) International PhD Program. This project has received funding from the European Union’s Horizon 2020 research and innovation programme under the Marie Sklodowska-Curie grant agreement No 665807, from Fondation Etchebès-Fondation France and Institut Carnot. We also would like to thank the Amgen Foundation and program for supporting JB and SGH.

## Supporting information

**S1 Fig. Increase of the *ank-katE* operon expression upon exposure to sublethal dose of H**_**2**_**O**_**2**_.

(A) Schematic representation of the *ank-katE* locus. The DNA fragments amplified by the PCR in (B) are designated with a bar and their corresponding size are indicated in base pairs in parenthesis.

(B) Electrophoresis gels of the PCR-amplified DNA fragments designated in (A) from genomic DNA (gDNA) or from mRNA before (RNA) or after a reverse transcriptase reaction (cDNA). DNA ladder fragment sizes are indicated at left of the gels.(

(C) Gene expression was measured by RT-qPCR reactions in WT *L. interrogans* exposed in the absence (white bars) or presence of 10 *μ*M H_2_O_2_ for 30 min (black bars) or 2h (dashed bars). Data are mean and SD of three independent experiments. ***, p-value<0.0001 by two-way Anova analysis.

**S2 Fig. Increase of *ahpC* and *ccp* expressions upon exposure to sublethal dose of H**_**2**_**O**_**2**_.

*L. interrogans* WT cells were cultivated until exponential phase and on exposed in the absence (white bars) or presence of 10 *μ*M H_2_O_2_ for 30 min (black bars) or for 2h (dashed bars). mRNAs were purified and cDNAs were subsequently prepared. *AhpC* (A), *sufB* (A), LIMLP_02790 (B) and *ccp* (B) expression was measured by RT-qPCR using *flaB* (LIMLP_09410) as reference gene and the data were normalized with untreated samples. Data are mean and SD of three independent experiments. ***, p-value<0.0001 by two-way Anova analysis.

**S3 Fig. Increase of the heme biosynthesis gene expression upon exposure to sublethal dose of H**_**2**_**O**_**2**_.

(A) Schematic representation of the heme cluster locus. The DNA fragments amplified by the PCR in (B) are designated with a bar and their corresponding size is indicated in base pairs in parenthesis.

(B) Electrophoresis gels of the PCR-amplified DNA fragments designated in (A) from genomic DNA (gDNA) or from mRNA before (RNA) or after a reverse transcriptase reaction (cDNA). DNA ladder fragment sizes are indicated at left of the gels.

(C) Gene expression was measured by RT-qPCR reactions in WT *L. interrogans* exposed in the absence (white bars) or presence of 10 *μ*M H_2_O_2_ for 30 min (black bars) or 2h (dashed bars). Data are mean and SD of three independent experiments. ****, p-value<0.0001; **, p-value<0.005; *, p-value<0.05 by two-way Anova analysis.

**S4 Fig. Increase of the LIMLP_17860-65 operon expression upon exposure to sublethal dose of H**_**2**_**O**_**2**_.

(A) Schematic representation of the locus of genes coding for a heme oxygenase and a permease. The DNA fragments amplified by the PCR in (B) are designated with a bar and their corresponding size is indicated in base pairs in parenthesis.

(B) Electrophoresis gels of the PCR-amplified DNA fragments designated in (A) from genomic DNA (gDNA) or from mRNA before (RNA) or after a reverse transcriptase reaction (cDNA). DNA ladder fragment sizes are indicated at left of the gels.

(C) Gene expression was measured by RT-qPCR reactions in WT *L. interrogans* exposed in the absence (white bars) or presence of 10 *μ*M H_2_O_2_ for 30 min (black bars) or 2h (dashed bars). Data are mean and SD of three independent experiments. *, p-value<0.05 by two-way Anova analysis.

**S5 Fig. Low binding of PerR with the PerR-controlled peroxidase locus.**

Chromatin immunoprecipitation was performed on *L. interrogans* WT and perR (M776) mutant strains in the presence or absence of the anti-PerR antibody. Co-immunoprecipitated DNA fragments located in the *ank-katE* operon locus (A), in the *ccp* locus (B) and in the *ahpC* locus (C) were amplified by qPCR. The location of amplified fragments is indicated below the schematic representation of their respective loci. The number of nucleotides between different ORFs is indicated in italic. Data are represented as fold enrichments.

**S6 Fig. Operon organization of the PerR-controlled TonB-dependent transport system locus.**

(A) Schematic representation of the locus of genes coding for a heme oxygenase and a permease. The DNA fragments amplified by the PCR in (B) are designated with a bar and their corresponding size is indicated in base pairs in parenthesis.

(B) Electrophoresis gels of the PCR-amplified DNA fragments designated in (A) from genomic DNA (gDNA) or from mRNA before (RNA) or after a reverse transcriptase reaction (cDNA). DNA ladder fragment sizes are indicated at left of the gels.

**S7 Fig. Low binding of PerR with the PerR-controlled TonB-dependent transport system locus.**

Chromatin immunoprecipitation was performed on *L. interrogans* WT and perR (M776) mutant strains in the presence or absence of the anti-PerR antibody. Co-immunoprecipitated DNA fragments located in the locus encoding a TonB-dependent transporter system were amplified by qPCR. The location of amplified fragments is indicated below the schematic representation of the respective locus. The number of nucleotides between different ORFs is indicated in italic. Data are represented as fold enrichments.

**S8 Fig. Operon organization of PerR-controlled *vicKR*.**

(A) Schematic representation of the locus of genes coding for the histidine kinase VicK and the response regulator VicR. The DNA fragments amplified by the PCR in (B) are designated with a bar and their corresponding size is indicated in base pairs in parenthesis.

(B) Electrophoresis gels of the PCR-amplified DNA fragments designated in (A) from genomic DNA (gDNA) or from mRNA before (RNA) or after a reverse transcriptase reaction (cDNA). DNA ladder fragment sizes are indicated at left of the gels.

