## Supplementary tables for "The adaptive transcriptional response of pathogenic *Leptospira* to peroxide reveals new defenses against infection-related oxidative stress": S1Table_ZavalaAlvarado.docx

| **ORF Id^a^** | **Gene** | **Fonction** | **Log_2_FC** | **Adjusted**  **p-value** | **RT-qPCR^b^** |
| --- | --- | --- | --- | --- | --- |
| ***Miscellaneous*** |  |  |  |  |  |
| LIMLP_05110 (LIC11058/ LA3017) |  | Lipoprotein LemA | 3.449* | 2.87e-44 |  |
| LIMLP_07150 (LIC11467/LA2498) | *ats1* | Chromosome condensation regulator RCC1 | 4.959* | 9.64e-47 | 21.851 |
| LIMLP_14170 (LIC10713/LA3469) | *irpA*/*lruB* | Peptidase M75/Imelysin/LruB | 2.822* | 4.04e-144 | 3.759 |
| LIMLP_14180 (LIC10711/LA3471) |  | Peptidase M75/Imelysin | 1.411 | 1.11e-16 | 1.663 |
| LIMLP_14465 (LIC10657/LA3540) | *sph* | Sphingomyelinase C | 2.044 | 4.62e-13 |  |
| LIMLP_18620 (LIC20152/LB192) |  | HmuY protein | 2.120* | 2.98e-38 | 3.153 |
| LIMLP_18625 (LIC20153/LB194) |  | Lipoprotein | 2.151* | 3.04e-84 | 2.705 |
| ***Regulators/signaling*** |  |  |  |  |  |
| LIMLP_04775 (LIC10996/LA3104) | *rtn* | Cyclic diguanylate phosphodiesterase/histidine kinase | 1.898 | 4.51e-37 |  |
| LIMLP_05055 (LIC11048/LA3033) |  | MolR transcriptional regulator | 1.119 | 3.55e-06 |  |
| LIMLP_05565 (LIC11146/LA2907) |  | DeoR transcriptional regulator | 2.389 | 7.31e-11 |  |
| LIMLP_05620 (LIC11158/LA2887) |  | Fur transcriptional regulator | 2.207* | 4.96e-16 |  |
| LIMLP_10155 (LIC12034/LA1857) | *perR* | Fur transcriptional regulator | 3.566*^#^ | 1.17e-83 | 6.105 |
| LIMLP_10945 (LIC12206/LA1576) |  | MarR EPS-associated transcriptional regulator | 2.301* | 4.16e-35 |  |
| LIMLP_11440 (LIC12305/LA1447) |  | LexA repressor | 2.402*^#^ | 1.64e-51 |  |
| LIMLP_12430 (LIC12490/LA1205) | *rpoE* | ECF sigma factor | 1.112 | 1.58e-09 |  |
| LIMLP_12515 (LIC12504/LA1186) |  | TCS response regulator CheY | 1.582 | 2.84e-18 |  |
| LIMLP_12520 (LIC12505/LA1185) |  | TCS response regulator | 1.282 | 4.47e-12 |  |
| LIMLP_14415 (LIC10666/LA3531) |  | ArsR transcriptional regulator | 1.65 | 2.04e-11 |  |
| LIMLP_15105 (LIC10525/LA3703) | *hrcA* | Heat-inducible repressor HrcA | 3.591*^#^ | 4.85e-24 | 6.768 |
| LIMLP_16805 (LIC13285/LA4122) | *rpoE* | ECF sigma factor | 1.285 | 4.85e-05 |  |
| ***Oxidative stress and redox-related*** |  |  |  |  |  |
| LIMLP_02795 (Lic12927/LA0666) | *ccp* | Cytochrome C peroxidase | 5.824*^#^ | 3.42e-218 | 41.68 |
| LIMLP_05955 (Lic11219/LA2809) | *ahpC* | Peroxiredoxin/alkylperoxiredoxin reductase | 4.007*^#^ | 1.41e-213 | 8.57 |
| LIMLP_05960 (Lic11220/LA2808) | *sufB* | ABC transporter permease | 2.234*^#^ | 4.81e-45 | 3.55 |
| LIMLP_07145 (LIC11466/LA2499) |  | Thiol oxidoreductase | 2.236^#^ | 1.50e-17 | 1.256 |
| LIMLP_07165 (LIC11470/LA2494) | *trxB* | Thioredoxin-disulfide reductase TrxB | 1.91^#^ | 9.98e-18 | 2.289 |
| LIMLP_08985 (LIC11810/LA2108) |  | Glutathione S-transferase | 1.23 | 3.66e-08 |  |
| LIMLP_10145 (LIC12032/LA1859) | *katE* | Catalase | 2.763*^#^ | 2.80e-90 | 6.086 |
| LIMLP_10150 (LIC12033/LA1858) |  | Ankyrin repeat-containing protein | 2.701*^#^ | 1.06e-89 | 2.424 |
| LIMLP_11965 (LIC12404/LA1321) | *dsbD* | Disulfide interchange protein | 1.474^#^ | 4.18e-18 |  |
| LIMLP_13670 (LIC10807-LEPIC0823/LA3356) | *yfcG*/*gst* | Glutathione S-transferase | 1.764 | 1.43e-28 | 2.305 |
| LIMLP_14175 (LIC10712/LA3470) |  | Thiol oxidoreductase | 1.802* | 1.02e-40 | 2.132 |
| LIMLP_14715 (LIC10606/LA3598) | *dps* | Ferritin/DNA-binding stress protein Dps | 1.095 | 8.91e-10 | 1.497 |
| LIMLP_18310 (LIC20093/LB117) | *ygaF*/*bcp* | Bacterioferritin comigratory protein/peroxiredoxin | 1.177 | 7.73e-11 | 1.353 |
| LIMLP_18595 (LIC20148/ LB186) | *pbsa* | Heme oxygenase | 1.179 | 5.57e-05 | 1.360 |
| LIMLP_18600 (LIC20149/ LB187) |  | Permease of the Major Facilitator Superfamily | 1.070 | 2.84e-04 |  |
| ***Chaperones*** |  |  |  |  |  |
| LIMLP_06540 (LIC11335/LA2655) | *groEL* | Molecular chaperone GroEL | 3.355*^#^ | 6.72e-35 | 6.936 |
| LIMLP_06545 (LIC11336/LA2654) | *groES* | Molecular chaperone GroES | 3.328*^#^ | 3.21e-33 | 4.862 |
| LIMLP_10060 (LIC12017/LA1879) | *clpB* | Disaggregating chaperone ClpB | 2.111*^#^ | 1.23e-15 | 2.487 |
| LIMLP_10970 (LIC12210/LA1564) | *ibpA* | Small heat shock protein Hsp20 | 6.788*^#^ | 1.33e-184 | 69.605 |
| LIMLP_10975 (LIC12211LA1563) | *hsp15* | Small heat shock protein Hsp15 | 6.589*^#^ | 1.45e-238 | 56.431 |
| LIMLP_15110 (LIC10525/LA3704) | *grpE* | GrpE | 3.610*^#^ | 4.47e-27 | 8.929 |
| LIMLP_15115 (LIC10524/LA3705) | *dnaK* | Molecular chaperone DnaK | 3.353*^#^ | 3.46e-31 | 6.667 |
| LIMLP_15120 (LIC10523/LA3706) | *dnaJ* | Molecular chaperone DnaJ | 2.611*^#^ | 4.81e-45 | 2.619 |
| ***DNA repair/SOS response*** |  |  |  |  |  |
| LIMLP_02170 (LIC13052/LA0503) | *dinP* | DNA polymerase IV/DNA damage inducible protein | 2.325* | 2.03e-36 |  |
| LIMLP_07780 (LIC11596/LA2351) |  | DNA mismatch repair protein MutS | 1.019^#^ | 1.19e-07 | 1.134 |
| LIMLP_07915 (LIC11620/LA2321) | *recN* | DNA repair protein RecN | 5.028*^#^ | 0.00 | 17.490 |
| LIMLP_08665 (LIC11745/LA2179) | *recA* | Recombinase RecA | 2.652* | 1.59e-58 |  |
| LIMLP_10880 (LIC12191/LA1589) |  | Mutator protein MutT/nudix hydrolase | 1.255 | 4.42e-04 |  |
| LIMLP_11400 (LIC12297/LA1456) |  | DNA repair protein RadC | 3.459* | 1.13e-167 |  |
| LIMLP_16520 (LIC10252/LA0294) |  | DNA repair exonuclease | 3.830*^#^ | 6.93e-63 | 6.113 |
| LIMLP_16525 (LIC10251/LA0293) |  | DNA repair Rad50 ATPase | 2.960*^#^ | 1.32e-86 | 2.899 |
| ***Transporter*** |  |  |  |  |  |
| LIMLP_04310 (LIC10902/LA3233) | *fecR* | Iron dicitrate transport regulator FecR | 1.594 | 3.70e-20 |  |
| LIMLP_07920 (LIC11621/LA2320) |  | Biopolymer transporter ExbB/TolQ | 2.080* | 1.47e-85 |  |
| LIMLP_07925 (LIC11622/LA2319) |  | Biopolymer transporter ExbD/Tol | 1.167 | 9.61e-21 |  |
| LIMLP_08410 (LIC11694/LA2242) |  | TonB-dependent receptor | 2.616* | 2.59e-20 |  |
| LIMLP_11395 (LIC12296/LA1457) |  | ABC transporter permease | 1.617* | 1.29e-31 |  |
| LIMLP_14160 (LIC10714/LA3468) | *fecA* | TonB-dependent receptor | 2.178*^#^ | 2.17e-83 | 2.743 |
| LIMLP_15535 (LIC10441/LA3806) | *amtB* | Ammonium transporter | 4.100* | 3.88e-50 |  |
| ***Prophage-related*** |  |  |  |  |  |
| LIMLP_00895 (LEPIC0178/LA0196) |  | Hypothetical | 3.661* | 4.81e-45 |  |
| LIMLP_04475 (LIC10401) |  | Hypothetical/bacteriophage related fragment | 1.278 | 2.91e-04 |  |
| LIMLP_04480 (no ortholog) |  | Hypothetical | 1.882 | 4.37e-09 |  |
| LIMLP_13010 (LIC12600/LA1067) |  | Hypothetical | 1.501 | 1.70e-07 |  |
| LIMLP_13015 (LIC12601/LA1066) |  | Hypothetical | 1.232 | 5.76e-05 |  |
| LIMLP_13020 (LIC12602/LA1065) |  | Hypothetical | 1.320 | 5.07e-06 |  |
| LIMLP_19610 (LA1831-33) |  | Phage replication protein | 1.176 | 2.26e-06 |  |
| ***Hypothetical*** |  |  |  |  |  |
| LIMLP_05115 (LIC11059/LA3016) |  | Hypothetical | 6.384* | 2.21e-108 |  |
| LIMLP_05120 (LEPIC1091/LA3015) |  | Hypothetical | 6.191* | 2.91e-04 |  |
| LIMLP_05555 (LIC11145/LA2909) |  | Hypothetical | 1.882 | 4.37e-09 |  |
| LIMLP_05560 (LIC11145/LA2908) |  | Hypothetical | 1.501 | 1.70e-07 |  |
| LIMLP_08415 (LIC11695/LA2241) |  | Hypothetical | 1.232 | 5.76e-05 |  |
| LIMLP_09650 (LIC11935/LA1968) |  | Hypothetical | 1.320 | 5.07e-06 |  |
| LIMLP_10275 (LIC12077/LA1725) |  | Phage replication protein | 1.176 | 2.26e-06 |  |
| LIMLP_11405 (LIC12298/LA1455) |  | Hypothetical | 6.384* | 2.21e-108 |  |
| LIMLP_12785 (LIC12555/LA1125) |  | Hypothetical | 6.191* | 2.91e-04 |  |
| LIMLP_13145 (LIC12628/LA1033) |  | Hypothetical | 1.882 | 4.37e-09 |  |
| LIMLP_13765 (LIC10790/LA3377) |  | Hypothetical | 1.501 | 1.70e-07 |  |
| LIMLP_17835 (LIC20007/LB009) |  | Hypothetical | 1.232 | 5.76e-05 |  |

**S3 Table: Selected up-regulated genes upon exposure to lethal doses of H_2_O_2_.**

Genes up-regulated upon 1 hour exposure to 1 mM H_2_O_2_.

^a^ Gene numeration is according to Satou et al. (2015).

^b^ Fold change (WT vs WT exposed 60 min to 1 mM H_2_O_2_) obtained by RT-qPCR experiment

* Genes significantly up-and down-regulated by Volcano analysis (Log_2_FC cutoff of 2 and p-value cutoff of 0.005).
