## Supplementary tables for "The adaptive transcriptional response of pathogenic *Leptospira* to peroxide reveals new defenses against infection-related oxidative stress": S2Table_ZavalaAlvarado.docx

| **ORF Id^a^** | **Gene** | **Function** | **Log_2_FC** | **Adjusted p-value** |
| --- | --- | --- | --- | --- |
| ***Protein synthesis/secretion*** |  |  |  |  |
| LIMLP_00815 (LIC10156/LA0177) | *yidD* | Membrane protein insertion effector | -1.490 | 2.52e-11 |
| LIMLP_00820 (LIC10157/LA0178) | *yidC* | Insertase | -1.357* | 1.98e-09 |
| LIMLP_03095 (LIC12870/LA0742) | *rplB* | 50S ribosomal protein L2 | -1.026 | 3.40e-03 |
| LIMLP_03100 (LIC12869/LA0743) | *rpsS* | 30S ribosomal protein L19 | -1.073 | 1.71e-03 |
| LIMLP_03105 (LIC12868/LA0744) | *rplV* | 50S ribosomal protein L22 | -1.263 | 9.62e-04 |
| LIMLP_03110 (LIC12867/LA0745) | *rpsC* | 30S ribosomal protein S3 | -1.176 | 1.45e-03 |
| LIMLP_03115 (LIC12866/LA0746) | *rplP* | 50S ribosomal protein L16 | -1.197 | 1.34e-03 |
| LIMLP_03120 (LIC12865/LA0747) | *rpmC* | 50S ribosomal protein L29 | -1.313 | 6.53e-04 |
| LIMLP_03125 (LIC12864/LA0748) | *rpsQ* | 30S ribosomal protein S17 | -1.388 | 6.41e-04 |
| LIMLP_03130 (LIC12863/LA0749) | *rplN* | 50S ribosomal protein L14 | -1.403 | 3.60e-04 |
| LIMLP_03135 (LIC12862/LA0750) | *rplX* | 50S ribosomal protein L24 | -1.323 | 4.32e-04 |
| LIMLP_03140 (LIC12861/LA0751) | *rplE* | 50S ribosomal protein L5 | -1.351 | 3.66e-04 |
| LIMLP_03150 (LIC12859/LA0753) | *rpsH* | 30S ribosomal protein S8 | -1.466 | 2.62e-04 |
| LIMLP_03155 (LIC12858/LA0754) | *rplF* | 50S ribosomal protein L6 | -1.496 | 1.95e-04 |
| LIMLP_03160 (LIC12857/LA0755) | *rplR* | 50S ribosomal protein L18 | -1.531 | 1.95e-04 |
| LIMLP_03165 (LIC12856/LA0756) | *rpsE* | 50S ribosomal protein L5 | -1.556 | 1.11e-04 |
| LIMLP_03170 (LIC12855/LA0757) | *rpmD* | 50S ribosomal protein L30 | -1.420 | 1.54e-04 |
| LIMLP_03175 (LIC12854/LA0758) | *rplO* | 50S ribosomal protein L15 | -1.322 | 1.54e-04 |
| LIMLP_03180 (LIC12853/LA0759) | *secY* | Translocon SecY subunit | -1.292 | 9.39e-05 |
| LIMLP_03190 (LIC12851/LA0761) | *infA* | translation initiation factor IF1 | -1.121 | 5.99e-04 |
| LIMLP_03195 (LIC12850) |  | 50S ribosomal L35 | -1.194 | 1.26e-03 |
| LIMLP_03200 (LIC12849/LA0762) | *rpsM* | 30S ribosomal L13 | -1.362 | 1.29e-04 |
| LIMLP_03205 (LIC12848/LA0763) | *rpsK* | 30S ribosomal S11 | -1.198 | 4.05e-04 |
| LIMLP_03210 (LIC12847/LA0764) | *rpsD* | 30S ribosomal S4 | -1.203 | 1.63e-04 |
| LIMLP_03215 (LIC12846/LA0765) | *rpoA* | RNA polymerase subunit alpha | -1.742 | 1.35e-06 |
| LIMLP_03220 (LIC12845/LA0766) | *yidC* | 50S ribosomal L17 | -1.693 | 2.64e-06 |
| LIMLP_07600 (LIC11557/LA2389) | *rimM* | Ribosome maturation/16S RNA processing | -1.260 | 4.98e-06 |
| LIMLP_12685 (LIC12537/LA1143) | *secF* | preprotein translocase SecF | -1.389 | 5.26e-07 |
| ***Cell respiration*** |  |  |  |  |
| LIMLP_03705 (LIC12752/LA0884) | *nuoN* | NADH quinone oxidoreductase subunit N | -1.390 | 2.08e-08 |
| LIMLP_03710 (LIC12751/LA0885) | *nuoN* | NADH quinone oxidoreductase subunit M | -1.478 | 2.90e-08 |
| LIMLP_03720 (LIC12749/LA0887) | *nuoL*/*nqo* | NADH quinone oxidoreductase subunit L12 | -1.161 | 2.30e-07 |
| LIMLP_03725 (LIC12748/LA0888) | *nuok* | NADH quinone oxidoreductase subunit K | -0.998 | 5.12e-06 |
| LIMLP_07965 (LIC11630/LA2309) | *fadD* | Long chain fatty acid CoA ligase/AMP binding protein | -1.891 | 7.93e-66 |
| LIMLP_10990 (LIC12214/LA1556) |  | Cytochrome C oxidase assembly factor SenC/SOC1 | -1.468 | 1.55e-20 |
| ***Metabolism*** |  |  |  |  |
| LIMLP_05260 (LIC11088/LA2974) | *maug* | Methylamine utilization protein | -2.296* | 4.11e-39 |
| LIMLP_06060 (LIC11240/LA2780) |  | ATP F0F1 synthase subunit δ | -1.283 | 7.23e-06 |
| LIMLP_06065 (LIC11241/LA2779) |  | ATP F0F1 synthase subunit α | -1.444 | 1.70e-06 |
| LIMLP_06070 (LIC11242/LA2778) |  | ATP F0F1 synthase subunit γ | -1.586 | 4.02e-07 |
| LIMLP_06075 (LIC11243/LA2776) |  | ATP F0F1 synthase subunit β | -1.670 | 2.78e-07 |
| LIMLP_06080 (LIC11244/LA2775) |  | ATP F0F1 synthase subunit ε | -1.434 | 1.14e-07 |
| LIMLP_18245 (LIC20080/LB103) | *ybgC*/*YbaW* | Acyl-CoA thioester hydrolase | -1.419 | 5.39e-08 |
| LIMLP_18455 (LIC20120/LB150) | *cobD* | Colabamin (VitB12) biosynthesis | -1.337 | 4.59e-07 |
| LIMLP_18460 (LIC20121/LB151) | *cobDQ* | Colabamin (VitB12) biosynthesis | -2.442* | 5.85e-21 |
| LIMLP_18465 (LIC20122/LB152) | *cobU* | Adenosylcobinamide | -2.536* | 5.65e-16 |
| LIMLP_18470 (LIC20123/LB153) |  | Adenosylcobinamide amidohydrolase | -2.180* | 1.15e-11 |
| LIMLP_18475 (LIC20124/LB154) | *cobB* | Cobyrinic acid a,c-diamide synthase | -2.085* | 5.60e-14 |
| LIMLP_18480 (LIC20125/LB155) | *cobA*/*btuR* | Cobyrinic acid a,c-diamide adenosyl transferase | -2.074* | 1.23e-11 |
| LIMLP_18485 (LIC20126/LB156) | *cobM*/*cbiF* | Precorrin-4 C11 methyltransferase | -2.099 | 3.98e-11 |
| LIMLP_18490 (LIC20127/LB157) | *cobJ*/*cbiH* | Precorrin-3B C17 methyltransferase | -2.441 | 1.58e-09 |
| LIMLP_18495 (LIC20128/LB158) | *cbiG* | Colabamin (VitB12) biosynthesis | -2.074 | 2.48e-08 |
| LIMLP_18500 (LIC20129/LB159) | *cobI*/*cobF* | Precorrin-2 C20 methyltransferase | -1.927 | 9.59e-09 |
| LIMLP_18505 (LIC20130/LB160) | *cobL*/*cbiET* | Precorrin-6Y C5, 15 methyltransferase | -1.935 | 2.08e-08 |
| LIMLP_18510 (LIC20131/LB161) | *cobH*/*cbiC* | Precorrin-8X methylmutase | -1.922 | 2.55e-07 |
| LIMLP_18515 (LIC20132/LB162) | *cbiD* | Cobalt precorrin 6A synthase | -1.659 | 6.93e-08 |
| LIMLP_18520 (LIC20133/LB163) |  | Oxidoreductase/FAD-binding flavodoxine reductase | -1.496 | 3.77e-09 |
| ***CRlSPR*** |  |  |  |  |
| LIMLP_02870 (LIC12914/LA0686) |  | CRISPR-associated protein Csh2 | -1.107 | 4.62e-13 |
| LIMLP_02875 (LIC12913/LA0687) |  | CRISPR-associated protein Cas8 | -1.285 | 3.45e-11 |
| LIMLP_02880 (LIC12912/LA0688) |  | CRISPR-associated protein Cas5 | -1.575 | 5.20e-06 |
| LIMLP_02885 (LIC12911-10/LA0689-90) |  | CRISPR-associated protein Cas3 | -1.212 | 8.70e-06 |
| ***Hypothetical*** |  |  |  |  |
| LIMLP_00510 (LIC10095/LA0107) |  | Hypothetical | -2.312* | 3.42e-14 |
| LIMLP_04180 (LIC12661/LA1000) |  | Hypothetical | -1.408 | 8.66e-13 |
| LIMLP_04220 (no ortholog) |  | Hypothetical | -1.414 | 3.49e-14 |
| LIMLP_04610 (LIC10963/LA3150) |  | Hypothetical | -1.370 | 6.59e-07 |
| LIMLP_05020 (LEPIC1072/LA3048) |  | Hypothetical | -2.452 | 3.70e-03 |
| LIMLP_05250 (LIC11086/LA2976) |  | Hypothetical | -1.676* | 2.18e-10 |
| LIMLP_05255 (LIC11087/LA2975) |  | Hypothetical | -2.446* | 3.95e-29 |
| LIMLP_05265 (LIC11089/LA2973) |  | Hypothetical | -2.100* | 3.03e-46 |
| LIMLP_07105 (LIC11458/LA2510) |  | Hypothetical | -1.328 | 4.56e-08 |
| LIMLP_07970 (LIC11631/LA2308) |  | Hypothetical | -1.327 | 1.79e-24 |
| LIMLP_11180 (LIC12253/LA1508) |  | Hypothetical | -1.574 | 1.16e-35 |
| LIMLP_11230 (LIC11262/LA1496) |  | Hypothetical | -1.612 | 4.88e-18 |
| LIMLP_11675 (LIC12343/LA1396) |  | Hypothetical | -1.359 | 9.78e-06 |
| LIMLP_11685 (LIC12345/LA1393) |  |  | -1.971 | 4.11e-12 |
| LIMLP_11780 (LIC12365/LA1366) |  | Hypothetical | -1.719 | 2.12e-11 |
| LIMLP_12590 (LIC12518/LA1168) |  | Hypothetical | -1.345 | 2.60e-09 |
| LIMLP_12910 (LIC12578/LA1097) |  | Hypothetical | -1.788 | 4.42e-07 |
| LIMLP_13720 (LIC10797/LA3368) |  | Hypothetical | -1.437 | 2.31e-08 |
| LIMLP_13725 (LIC10796/LA3369) |  | Hypothetical | -1.389 | 3.29e-09 |
| LIMLP_14190 (LIC10709/LEPIN3051) |  | Hypothetical | -2.128 | 5.58e-10 |
| LIMLP_14195 (LIC10708/LA3473) |  | Hypothetical | -1.702 | 9.80e-08 |
| LIMLP_14450 (LIC10660/LA3537) |  | Hypothetical | -1.701 | 2.14e-09 |
| LIMLP_15090 (LIC10529/LA3697) |  | Hypothetical | -1.407 | 1.94e-12 |
| LIMLP_15315 (LIC10980/LA3752) |  | Hypothetical | -1.444 | 5.39e-08 |
| LIMLP_15335 (LIC10476/LA3756) |  | Hypothetical | -1.545 | 2.84e-04 |
| LIMLP_15620 (no ortholog) |  | Hypothetical | -1.736 | 1.98e-04 |
| LIMLP_15715 (LIC10140/LA0470) |  | Hypothetical | -1.834 | 2.64e-12 |
| LIMLP_16170 (LEPIC0338/LA0371) |  | Hypothetical | -1.613 | 1.08e-11 |
| LIMLP_17425 (LIC12518/LA1168) |  | Hypothetical | -1.402 | 7.59e-06 |
| LIMLP_17465 (LA4271) |  | Hypothetical | -1.814 | 3.66e-04 |
| LIMLP_17470 (LIC13426/LA4280) |  | Hypothetical | -1.693 | 6.02e-04 |
| LIMLP_18675 (LIC20162/LB205) |  | Hypothetical | -1.372 | 1.28e-07 |
| LIMLP_18680 (LIC20163/LB207) |  | Hypothetical | -1.910 | 2.69e-12 |
| LIMLP_19115 (LIC20244/LB320) |  | Hypothetical | -1.324 | 2.41e-05 |

**S4 Table: Selected down-regulated genes upon exposure to lethal doses of H_2_O_2_.**

Genes down-regulated upon 1 hour exposure to 1 mM of H_2_O_2_.

^a^ Gene numeration is according to Satou et al. (2015). Corresponding genes of *L. interrogans* serovar lai strain 56601 and *L. interrogans* serovar Copenhageni Fiocruz strain L1-130 are indicated in parenthesis.

* Genes significantly down-regulated by Volcano analysis (Log_2_FC cutoff of 2 and p-value cutoff of 0.005).
