## Supplementary tables for "The adaptive transcriptional response of pathogenic *Leptospira* to peroxide reveals new defenses against infection-related oxidative stress": S3Table_ZavalaAlvarado.docx

| **Mutant name** | **ORF** | **Annotation** | **Position in genome^a^** | **Tn insertion site^b^** |
| --- | --- | --- | --- | --- |
| Man 69^c^ | LIMLP_10145 | Catalase (*katE*) | 2425442-2426887 | 2425862 |
| M776^d^ | LIMLP_10155 | PerR | 2427923-2428360 | 2427985 |
| Man 782^c^ | LIMLP_04245 | Biopolymer transport ExbD/TolR | 1029081-1029491 | 1029153 |
| Man 899^c^ | LIMLP_16720 | Two component sytem response regulator (*vicR*) | 3977080-3977781 | 3977570 |
| Man 1022^c^ | LIMLP_04270 | TonB-dependent receptor | 1032369-1035344 | 1033465 |
| Man 1368^c^ | LIMLP_05955 | Peroxiredoxin (*ahpC*) | 1458932-1459513 | 1459214 |
| Man 1448^c^ | LIMLP_16725 | Two component system histidine kinase (*vicK*) | 3977771-3979162 | 3978896 |

^a^ position according to *Leptospira interrogans* serovar Manilae UP-MMC-NIID-LP genome

^b^ nucleotide of the insertion site according to *Leptospira interrogans* serovar Manilae UP-MMC-NIID-LP genome

^c^ Mutants obtained by random insertion of a transposon (our laboratory)

^d^ Mutant obtained by random insertion of a transposon (Lo et al. (2010) Infect. Immun. Vol. 78, 4850-4859)

**S6 Table: Transposon mutants used in this study**.
