## Supplementary figures for "The adaptive transcriptional response of pathogenic *Leptospira* to peroxide reveals new defenses against infection-related oxidative stress": S1Figure_ZavalaAlvarado.pptx

### Slide 1
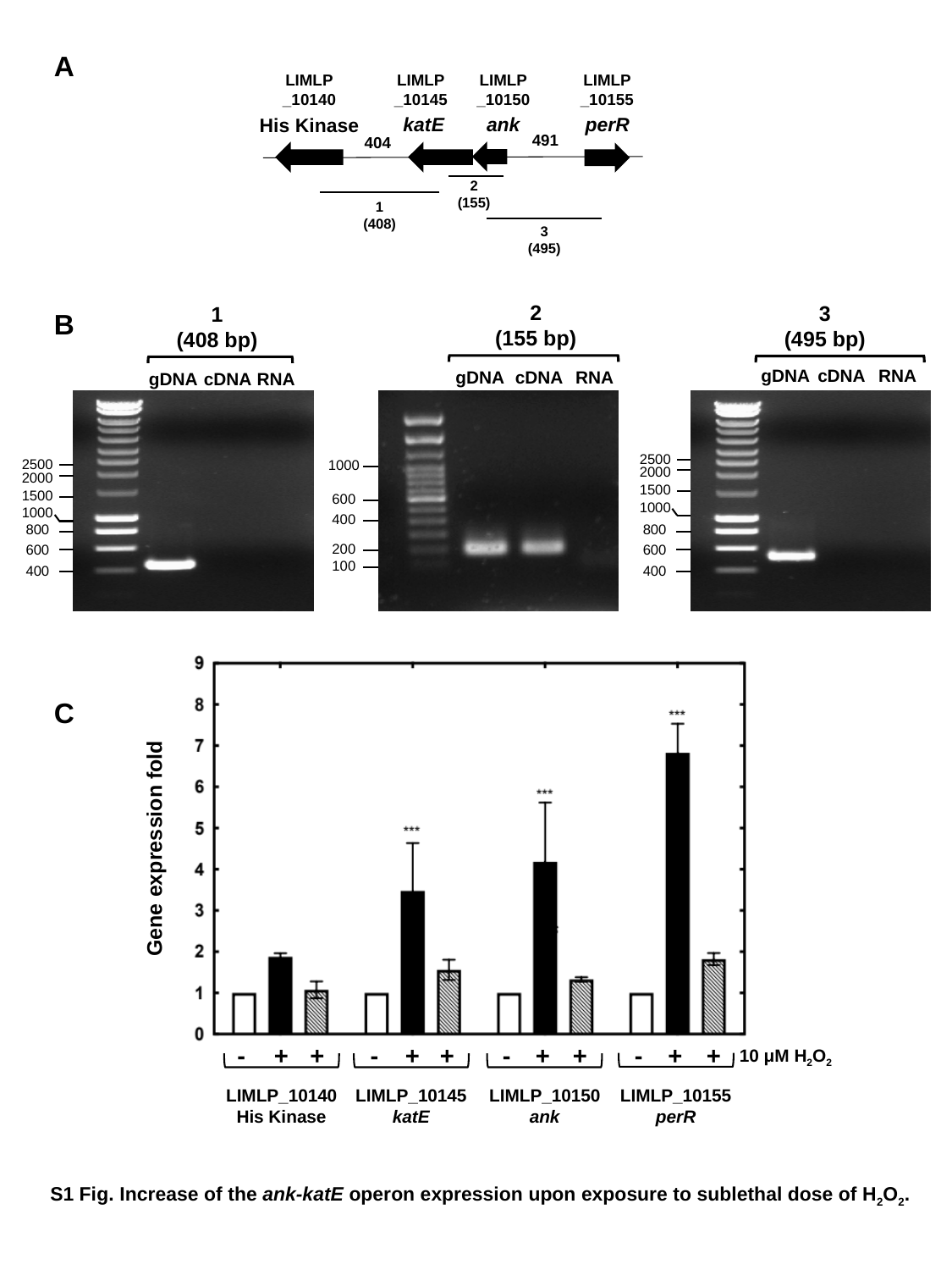

A
LIMLP
_10140
His Kinase
LIMLP
_10145
 katE
LIMLP
_10150
ank
LIMLP
_10155
perR
491
404
2
(155)
1
(408)
3
(495)
2
(155 bp)
gDNA
cDNA
RNA
1000
600
400
200
100
3
(495 bp)
gDNA
cDNA
RNA
800
600
400
1
(408 bp)
gDNA
cDNA
RNA
2500
2000
1500
1000
800
600
400
B
C
Gene expression fold
-
+
+
LIMLP_10140
His Kinase
-
+
+
LIMLP_10145
katE
-
+
+
LIMLP_10150
ank
-
+
+
LIMLP_10155
perR
10 μM H_2_O_2_
S1 Fig. Increase of the ank-katE operon expression upon exposure to sublethal dose of H_2_O_2_.
2500
2000
1500
1000
