## Supplementary figures for "The adaptive transcriptional response of pathogenic *Leptospira* to peroxide reveals new defenses against infection-related oxidative stress": S2Figure_ZavalaAlvarado.pptx

### Slide 1
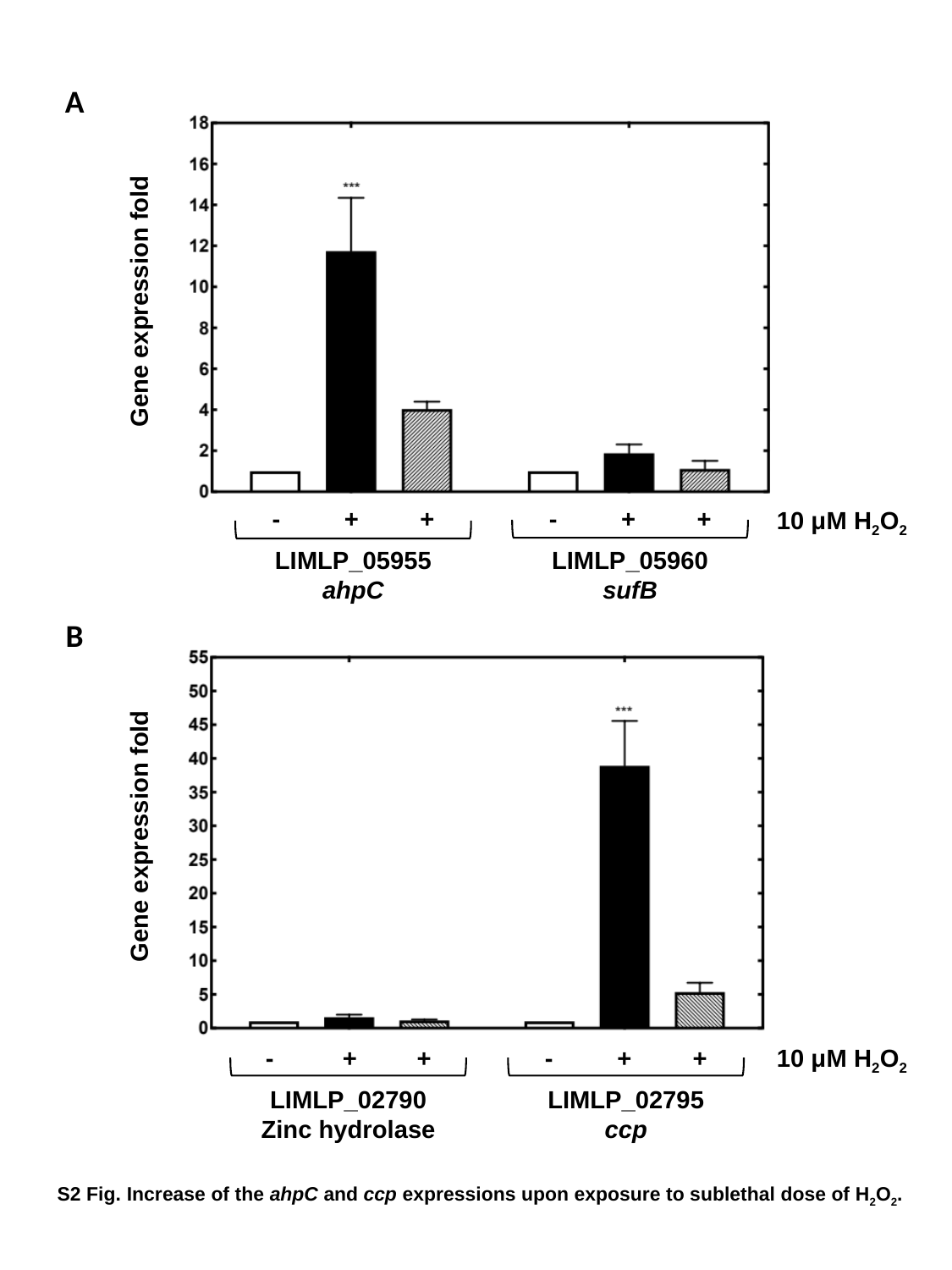

A
Gene expression fold
-
+
+
LIMLP_05955
ahpC
-
+
+
LIMLP_05960
sufB
10 μM H_2_O_2_
B
Gene expression fold
-
+
+
LIMLP_02790
Zinc hydrolase
-
+
+
LIMLP_02795
ccp
10 μM H_2_O_2_
S2 Fig. Increase of the ahpC and ccp expressions upon exposure to sublethal dose of H_2_O_2_.
