## Supplementary figures for "The adaptive transcriptional response of pathogenic *Leptospira* to peroxide reveals new defenses against infection-related oxidative stress": S3Figure_ZavalaAlvarado.pptx

### Slide 1
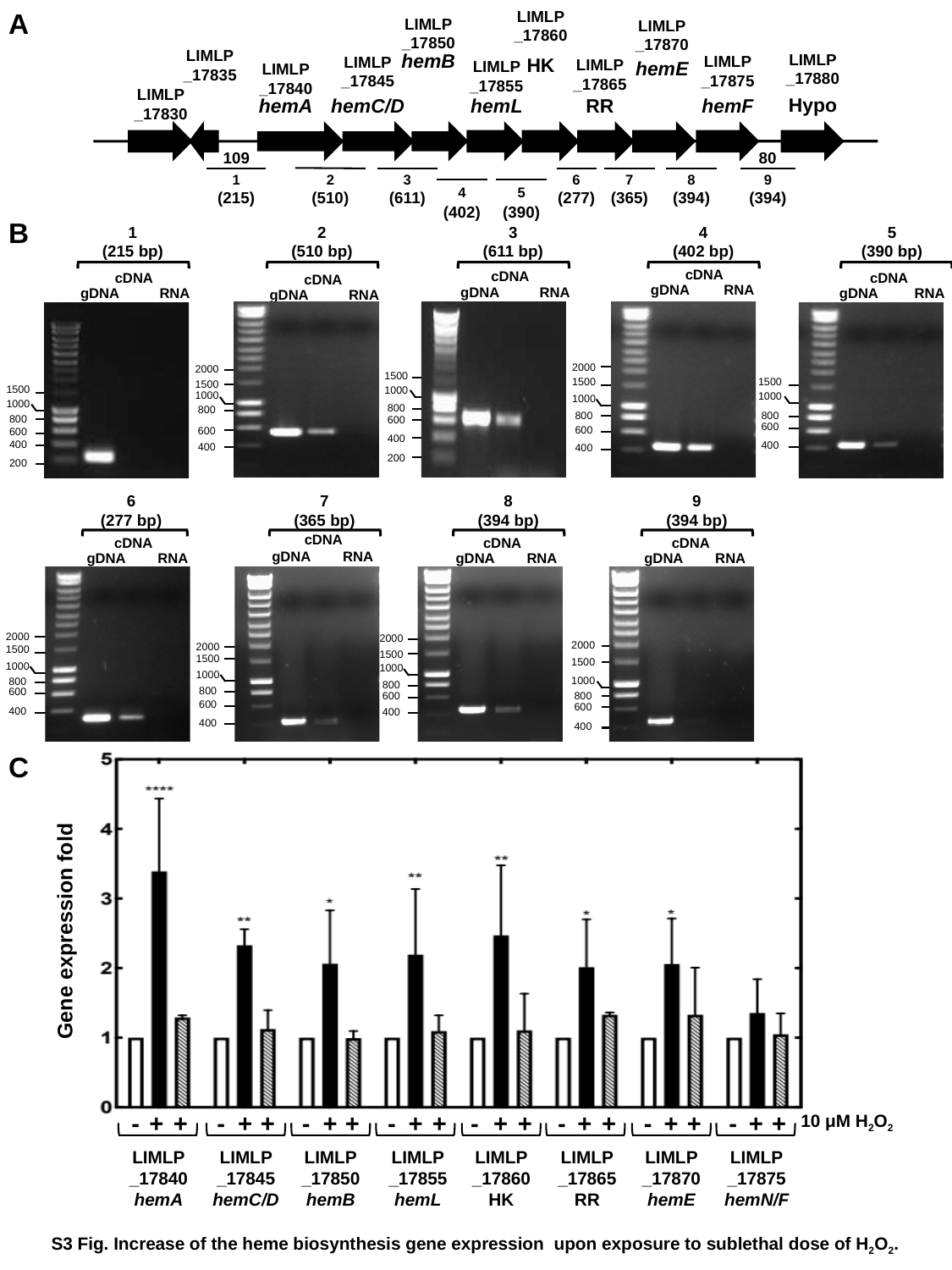

LIMLP
_17860
LIMLP
_17850
LIMLP
_17870
LIMLP
_17835
hemB
LIMLP
_17880
LIMLP
_17875
LIMLP
_17845
HK
LIMLP
_17865
hemE
LIMLP
_17855
LIMLP
_17840
LIMLP
_17830
Hypo
hemL
hemA
hemC/D
RR
hemF
109
80
2
(510)
1
(215)
3
(611)
6
7
8
9
4
(402)
5
(277)
(365)
(394)
(394)
(390)
A
B
1
(215 bp)
gDNA
RNA
1500
1000
800
600
400
200
cDNA
2
(510 bp)
cDNA
gDNA
RNA
2000
1500
1000
800
600
400
3
(611 bp)
cDNA
gDNA
RNA
1500
1000
800
600
400
200
4
(402 bp)
cDNA
gDNA
RNA
2000
1500
1000
800
600
400
5
(390 bp)
cDNA
gDNA
RNA
1500
1000
800
600
400
6
(277 bp)
cDNA
gDNA
RNA
2000
1500
1000
800
600
400
7
(365 bp)
cDNA
gDNA
RNA
2000
1500
1000
800
600
400
8
(394 bp)
cDNA
gDNA
RNA
2000
1500
1000
800
600
400
9
(394 bp)
cDNA
gDNA
RNA
2000
1500
1000
800
600
400
Gene expression fold
-
+
+
LIMLP
_17840
hemA
-
+
+
LIMLP
_17845
hemC/D
-
+
+
LIMLP
_17850
hemB
-
+
+
LIMLP
_17855
hemL
-
+
+
LIMLP
_17860
HK
-
+
+
LIMLP
_17865
RR
-
+
+
LIMLP
_17870
hemE
-
+
+
LIMLP
_17875
hemN/F
10 μM H_2_O_2_
C
S3 Fig. Increase of the heme biosynthesis gene expression upon exposure to sublethal dose of H_2_O_2_.
