## Supplementary figures for "The adaptive transcriptional response of pathogenic *Leptospira* to peroxide reveals new defenses against infection-related oxidative stress": S4Figure_ZavalaAlvarado.pptx

### Slide 1
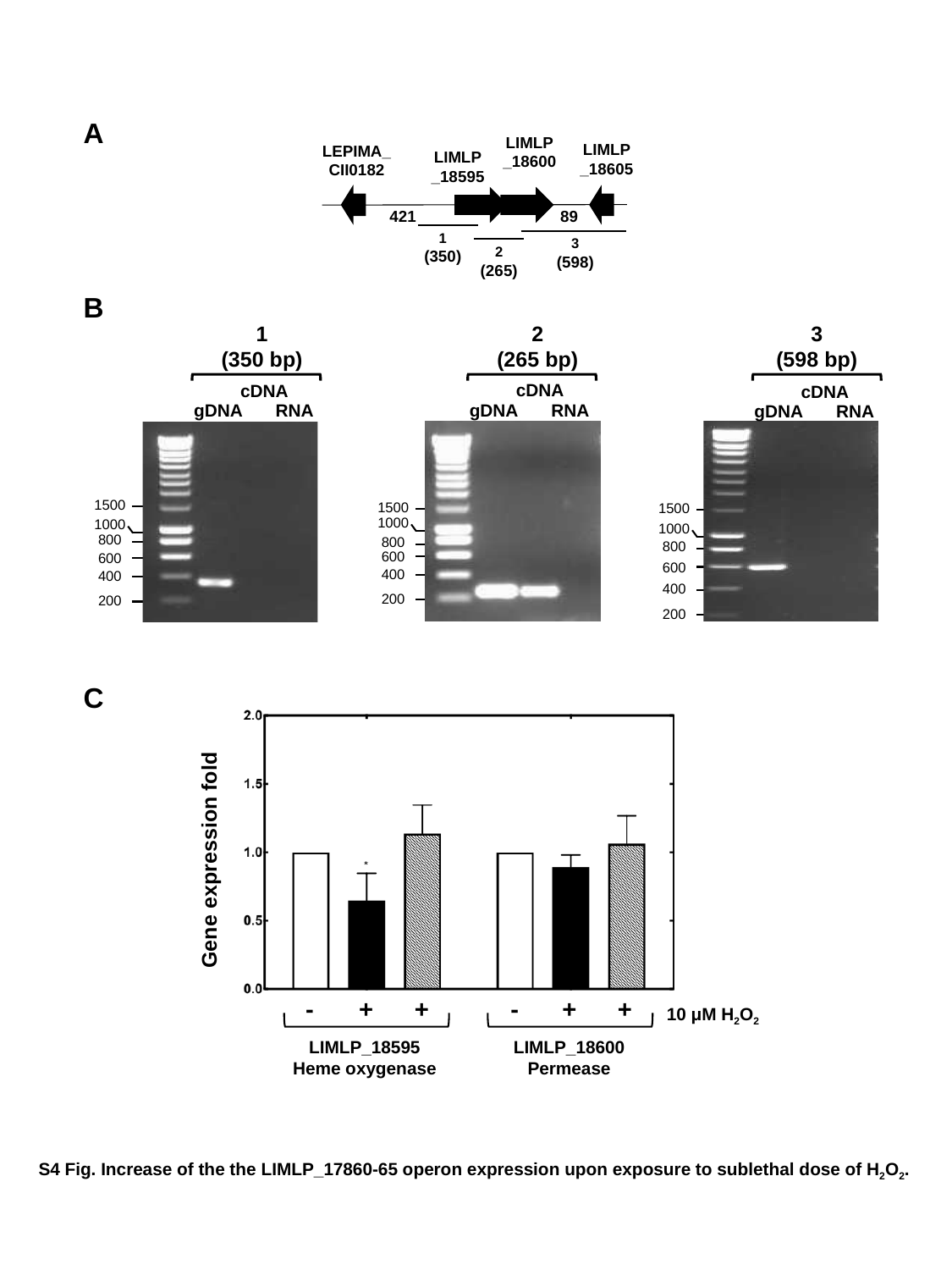

A
LIMLP
_18600
LIMLP
_18605
LEPIMA_CII0182
LIMLP
_18595
421
89
1
(350)
3
(598)
2
(265)
B
1
(350 bp)
cDNA
gDNA
RNA
1500
1000
800
600
400
200
2
(265 bp)
cDNA
gDNA
RNA
1500
1000
800
600
400
200
3
(598 bp)
cDNA
gDNA
RNA
1500
1000
800
600
400
200
C
Gene expression fold
-
+
+
LIMLP_18595
Heme oxygenase
-
+
+
LIMLP_18600
Permease
10 μM H_2_O_2_
S4 Fig. Increase of the the LIMLP_17860-65 operon expression upon exposure to sublethal dose of H_2_O_2_.
