## Supplementary figures for "The adaptive transcriptional response of pathogenic *Leptospira* to peroxide reveals new defenses against infection-related oxidative stress": S5Figure_ZavalaAlvarado.pptx

### Slide 1
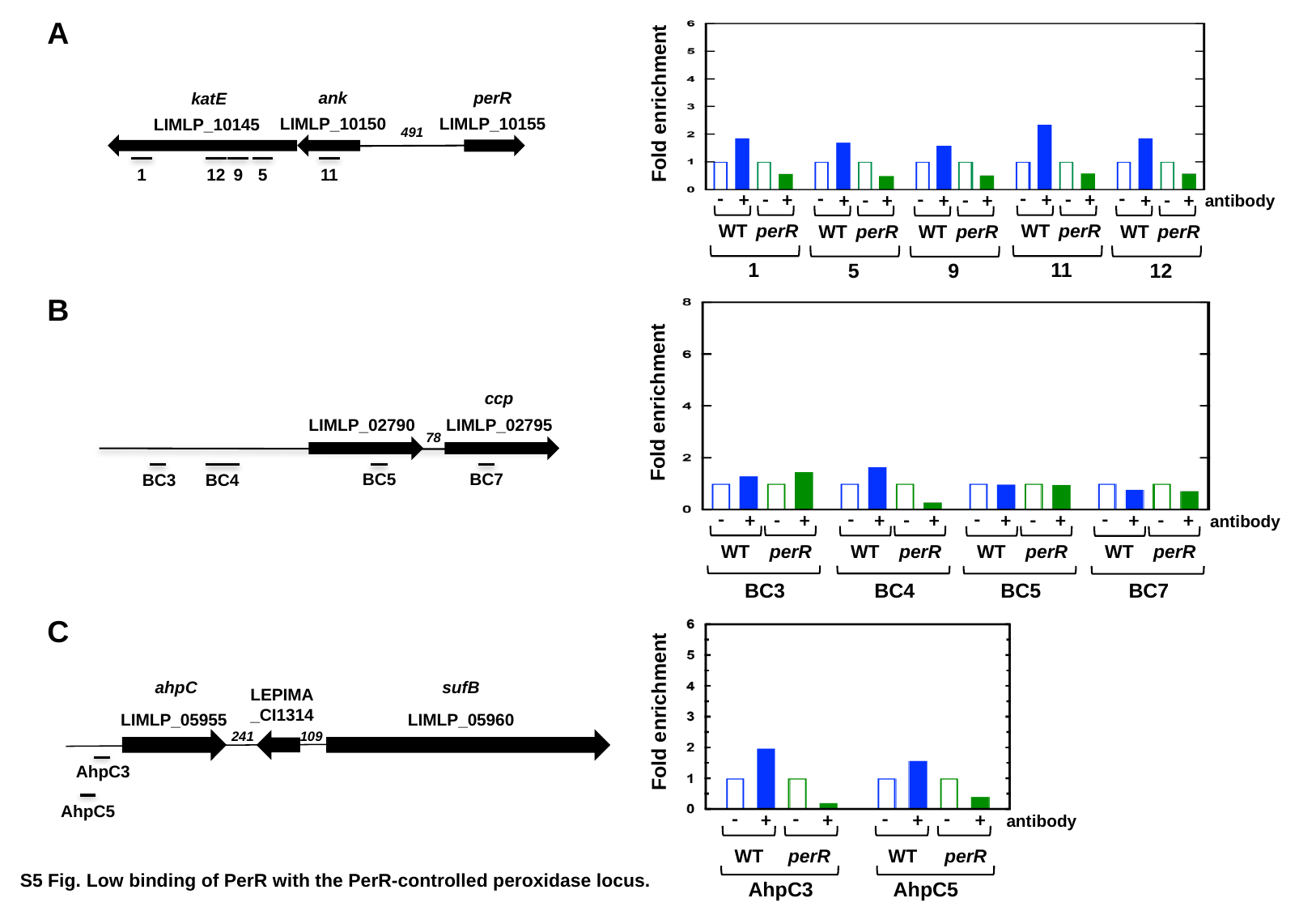

Fold enrichment
-
+
WT
-
+
perR
11
-
+
WT
-
+
perR
1
-
+
WT
-
+
perR
12
-
+
WT
-
+
perR
5
-
+
WT
-
+
perR
9
antibody
perR
LIMLP_10155
ank
LIMLP_10150
 katE
LIMLP_10145
11
1
12
9
5
491
A
B
Fold enrichment
-
+
WT
-
+
perR
BC3
-
+
WT
-
+
perR
BC4
-
+
WT
-
+
perR
BC5
-
+
WT
-
+
perR
BC7
antibody
ccp
LIMLP_02795
LIMLP_02790
BC3
BC4
BC5
BC7
78
Fold enrichment
-
+
perR
-
+
WT
AhpC5
-
+
perR
-
+
WT
AhpC3
antibody
 ahpC
LIMLP_05955
sufB
LIMLP_05960
LEPIMA
_CI1314
241
109
AhpC3
AhpC5
C
S5 Fig. Low binding of PerR with the PerR-controlled peroxidase locus.
