## Supplementary figures for "The adaptive transcriptional response of pathogenic *Leptospira* to peroxide reveals new defenses against infection-related oxidative stress": S6Figure_ZavalaAlvarado.pptx

### Slide 1
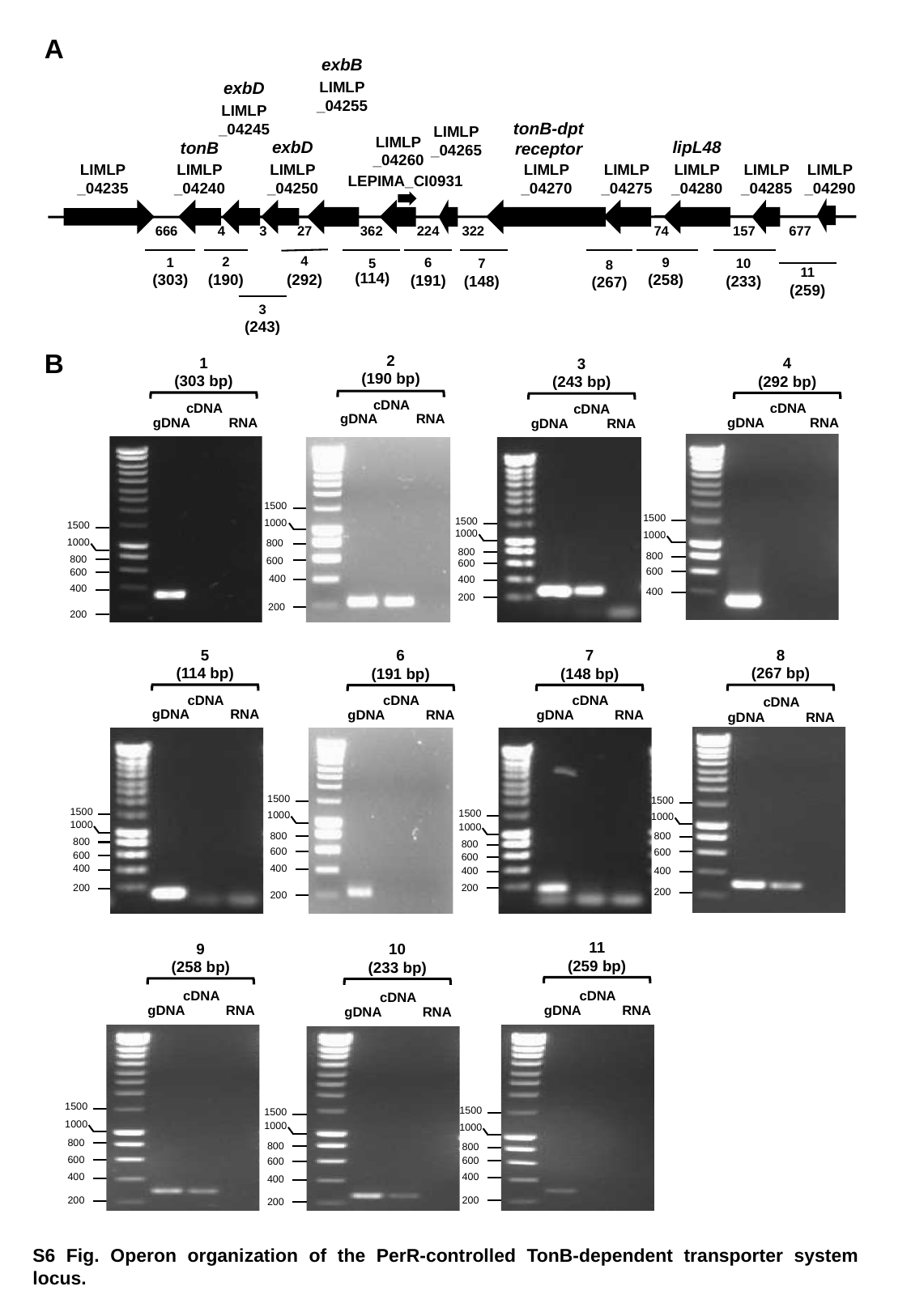

A
exbB
LIMLP
_04255
exbD
LIMLP
_04245
tonB-dpt receptor
lipL48
LIMLP
_04280
LIMLP
_04265
LIMLP
_04260
exbD
LIMLP
_04250
tonB
LIMLP
_04240
LIMLP
_04235
LIMLP
_04270
LIMLP
_04275
LIMLP
_04285
LIMLP
_04290
LEPIMA_CI0931
666
4
3
27
362
224
322
74
157
677
4
(292)
1
(303)
2
(190)
5
(114)
6
(191)
7
(148)
8
(267)
9
(258)
10
(233)
11
(259)
3
(243)
B
2
(190 bp)
cDNA
gDNA
RNA
1500
1000
800
400
200
600
1
(303 bp)
cDNA
gDNA
RNA
1500
1000
800
400
200
600
4
(292 bp)
cDNA
gDNA
RNA
1500
1000
800
400
600
3
(243 bp)
cDNA
gDNA
RNA
1500
1000
800
400
200
600
5
(114 bp)
cDNA
gDNA
RNA
1500
1000
800
400
200
600
8
(267 bp)
cDNA
gDNA
RNA
1500
1000
800
400
200
600
7
(148 bp)
cDNA
gDNA
RNA
1500
1000
800
400
200
600
6
(191 bp)
cDNA
gDNA
RNA
1500
1000
800
400
200
600
11
(259 bp)
cDNA
gDNA
RNA
1500
1000
800
400
200
600
9
(258 bp)
cDNA
gDNA
RNA
1500
1000
800
400
200
600
10
(233 bp)
cDNA
gDNA
RNA
1500
1000
800
400
200
600
S6 Fig. Operon organization of the PerR-controlled TonB-dependent transporter system locus.
