## Supplementary figures for "The adaptive transcriptional response of pathogenic *Leptospira* to peroxide reveals new defenses against infection-related oxidative stress": S7Figure_ZavalaAlvarado.pptx

### Slide 1
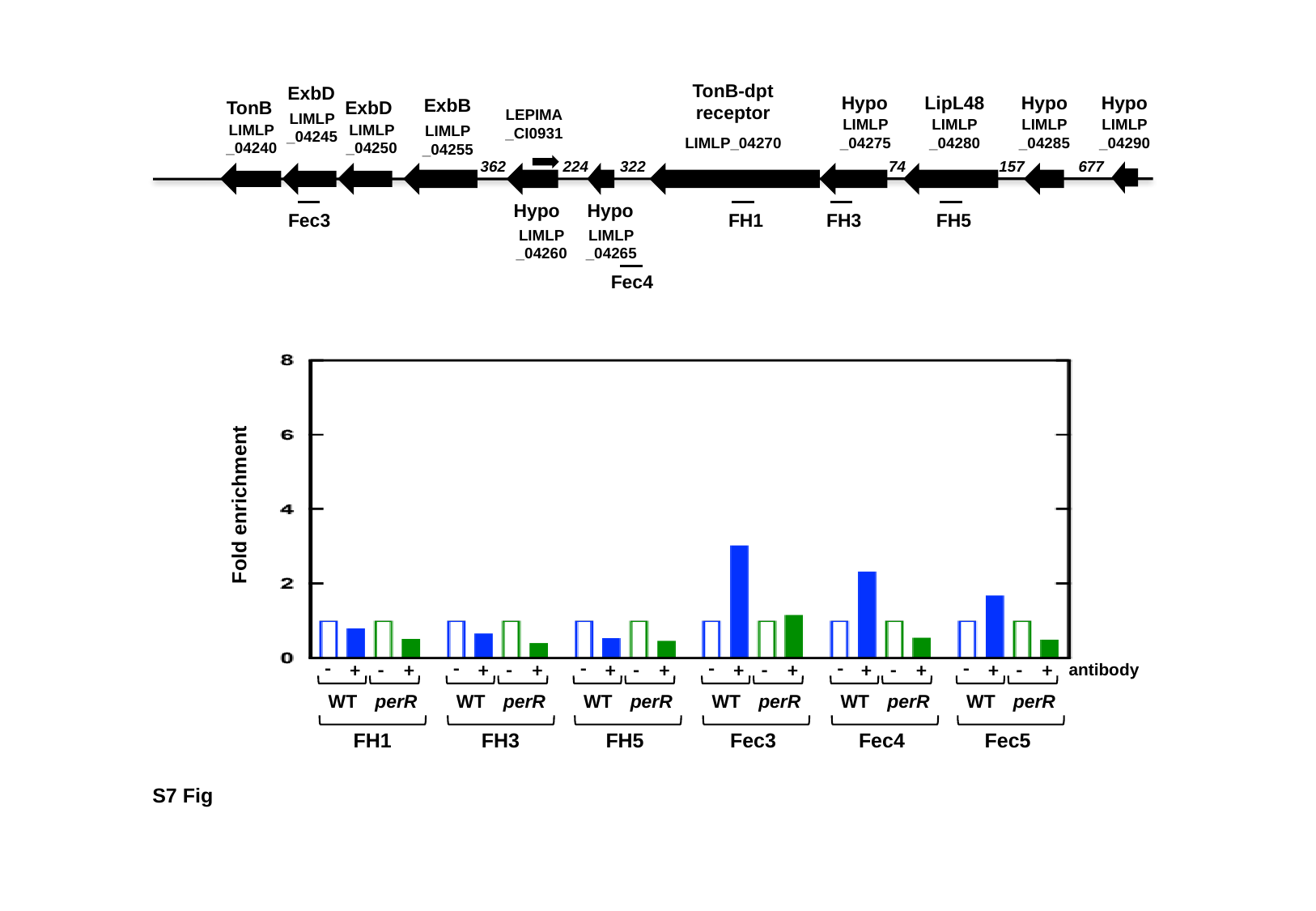

TonB-dpt
receptor
LIMLP_04270
224
322
74
157
677
362
ExbD
LIMLP
_04245
Hypo
LIMLP
_04275
LipL48
LIMLP
_04280
Hypo
LIMLP
_04285
Hypo
LIMLP
_04290
ExbB
LIMLP
_04255
TonB
LIMLP
_04240
ExbD
LIMLP
_04250
LEPIMA
_CI0931
Hypo
LIMLP
_04260
Hypo
LIMLP
_04265
Fec3
FH1
FH3
FH5
Fec4
Fold enrichment
-
+
WT
-
+
perR
FH1
-
+
WT
-
+
perR
FH3
-
+
WT
-
+
perR
FH5
-
+
WT
-
+
perR
Fec3
-
+
WT
-
+
perR
Fec4
-
+
WT
-
+
perR
Fec5
antibody
S7 Fig
