## Supplementary figures for "The adaptive transcriptional response of pathogenic *Leptospira* to peroxide reveals new defenses against infection-related oxidative stress": S8Figure_ZavalaAlvarado.pptx

### Slide 1
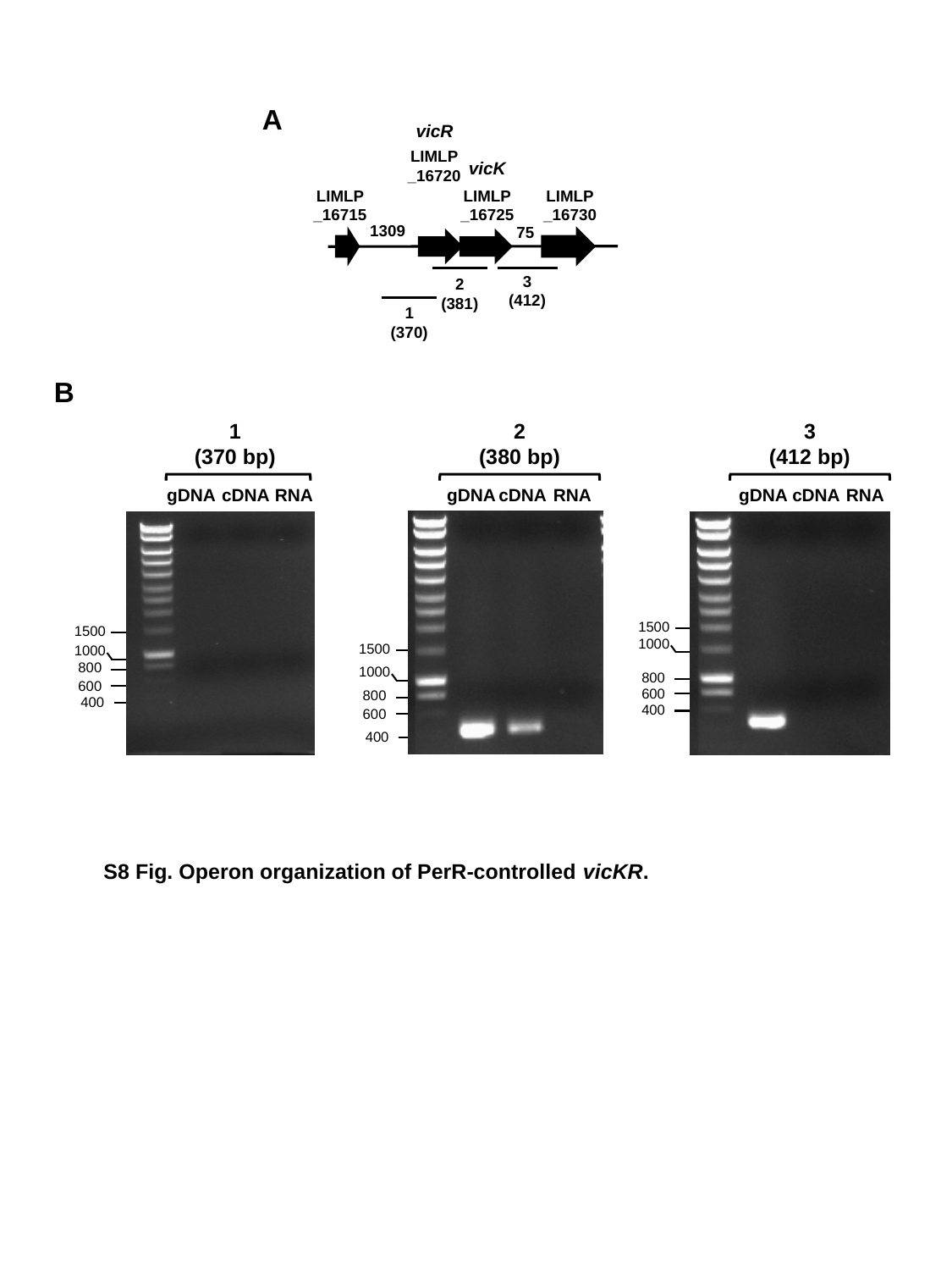

A
vicR
LIMLP
_16720
vicK
LIMLP
_16725
LIMLP
_16715
LIMLP
_16730
1309
75
2
(381)
3
(412)
1
(370)
B
1
(370 bp)
gDNA
cDNA
RNA
1500
1000
800
600
400
2
(380 bp)
gDNA
cDNA
RNA
1500
1000
800
600
400
3
(412 bp)
gDNA
cDNA
RNA
1500
1000
800
600
400
S8 Fig. Operon organization of PerR-controlled vicKR.
